# Specific neuroblast-derived signals control both cell migration and fate in the rostral migratory stream

**DOI:** 10.1101/2025.02.18.638163

**Authors:** Wei Zhou, James R. Munoz, Hsin-Yi Henry Ho, Kenji Hanamura, Matthew B. Dalva

## Abstract

Functional neuronal circuits require neuroblasts migrate to appropriate locations and then differentiate into neuronal subtypes. However, it remains unknown how neuroblasts in the subventricular zone (SVZ) are guided through the rostral migratory stream (RMS) to the olfactory bulb (OB). Here we define EphB2 as a neuroblast-derived cue that controls migration along the RMS and helps to determine cell fate. Within the RMS, EphB2 is expressed selectively in, kinase-active in, and required for the migration of neuroblasts. As neuroblasts enter the OB and differentiate, EphB kinase activity is down-regulated, and in the granule cell layer (GCL), EphB2 expression is down-regulated. Blocking EphB kinase activity or knocking down EphB2 results in defects in migration and premature cellular differentiation in the RMS. Unexpectedly, premature loss of EphB2 expression causes neuroblasts to stop migrating and differentiate into astrocyte-like cells. Thus, EphB2 kinase activity and expression are linked to migration and specification of neuroblast fate.

## Introduction

The subventricular zone (SVZ) of the lateral ventricle is the major site of neurogenesis in the developing and adult brain. Newly generated neuroblasts migrate along the wall of the lateral ventricle in close contact with one another [1-3] and then chain migrate through the astrocyte-ensheathed rostral migratory stream (RMS) to the olfactory bulb (OB). Cells that migrate into the OB must then migrate radially to reach the granule cell layer (GCL) and differentiate into inhibitory interneurons [4]. With migration and differentiation complete, new inhibitory interneurons integrate into the circuitry of the OB. These events are likely playing key functions in olfactory learning, and have been observed in young primates including humans [5, 6]. Moreover, olfactory dysfunction is found in schizophrenia and Alzheimer’s disease, with reduced olfactory bulb size and function associated with disease [7-10]. However, despite the importance of the migratory path these neurons must follow, the specific molecular cues guide migrating neuroblasts along the RMS to the OB and whether these cues regulate the decision of the neuroblasts to differentiate into mature cell types remain unresolved.

One important element of migration along the RMS are the ensheathing glial cells that surround neuroblasts along the RMS. Mutations that result in defective formation of the glial tube alters migration along the RMS, suggests that cell-to-cell interactions between migrating neuroblasts and glia may help to guide cells along the correct pathway [11-13]. Indeed, in the adult rodent brain SVZ and RMS axon guidance family molecules are expressed in glial cells surrounding the RMS and in migrating neuroblasts including the Ephs-ephrins, PSA-NCAM, Slit/Robo, and Sonic Hedgehog systems [13-15]. These guidance molecules are thought to regulate migration of neuroblasts along the RMS. For instance, Slit/Robo signaling between glial cells and neuroblasts may determine the speed of migration along the RMS, while PSA-NCAM signaling appears to control the width of the RMS [16, 17]. Yet the molecular cue or cues that enable the RMS ensheathing glial cells to keep migrating neuroblasts within the RMS are unknown.

Of the guidance molecules expressed in the RMS region, the Eph family receptor tyrosine kinases (RTKs) are attractive candidates. EphB RTKs are a family of five transmembrane signaling molecules. Binding of one of three transmembrane proteins known as ephrin-Bs (eB1-3) activates the EphB kinase. EphB signaling regulates events that require cell-cell contact throughout development and in the mature brain [18, 19]. In a variety of migrating cell types, EphB signaling regulates cell morphology and motility [18, 20-23]. In addition, infusion of ephrin-B2 or EphB2 ecto-domains, but not ephrin-B1, into the lateral ventricle of mice disrupts migration and increases proliferation of neuroblasts in the SVZ [24] and recent studies have linked EphAs to the regulation of migration within the SVZ and RMS [14]. The finding that ephrin-B1 ectodomains do disrupt proliferation is important as it indicates that proliferation is likely to be mediated by EphB4, which is the only EphB protein known to be activated by ephrin-B2 but not ephrin-B1. It is not known whether EphB proteins play a specific role in guiding neuroblasts through the RMS to the olfactory bulb.

Here we define the function of EphB2 and EphB signaling in neuroblast migration through the RMS and into the OB. EphB2 is selectively expressed and kinase-active in neuroblasts during tangential chain migration along the RMS. Once cells reach the OB, EphB kinase activity is down-regulated, and cells migrate radially into the granule cell layer (GCL). As neuroblasts undergo terminal differentiation, expression of EphB2 is lost. We propose that EphB kinase activity is required for chain migration, while maintenance of EphB2 expression along the RMS may regulate the cell fate of cells that are destined to become neurons.

Consistent with this model, *in vivo* blockade of EphB kinase activity using a chemo-genetic mouse model results in defective migration and drives cells to begin radial migration prematurely while still in the RMS. *In vivo* loss of EphB2 expression in single neuroblasts prevents cell migration and drives neuroblasts to undergo inappropriate differentiation. These data support a model where downregulation of EphB2 kinase activity is a signal for cells to migrate radially, and the loss of EphB2 expression leads to cell differentiation. Together these findings suggest that neuroblasts must complete their migration along the correct path to achieve appropriate cell fate.

## Results

### EphB2 is expressed and phosphorylated in chain migrating neuroblasts

EphB signaling can regulate the motility of a variety of migrating cell types, and EphBs and ephrin-Bs are expressed in the SVZ [14, 20-22, 24]. Therefore, we asked whether EphBs were also expressed in cells migrating from the SVZ to the OB within the RMS. Sections containing the RMS of adult mouse brain were stained with EphB1, EphB2, or EphB3 antibodies and markers for neuroblasts (doublecortin (DCX)) or astrocytes/type B cells (GFAP; Supplementary Figure 1, 2). The specificities of the EphB antibodies were confirmed by staining sections from EphB1-3 triple knockout mice (EphB TKO; EphB1^-/-^, EphB2^-/-^, EphB3^-/-^; Figure 1 – Figure supplement 1E-G). We find that EphB1 and EphB3 are expressed in the GFAP+ astrocytes that line the RMS (Supplementary Figure 1A, 1B and 1D, colocalization indicated by purple color, Supplementary Figure 2A and 2C). In contrast, EphB2 is expressed in chain migrating DCX+ neuroblasts (Supplementary Figure 1C).

**Figure 1.**
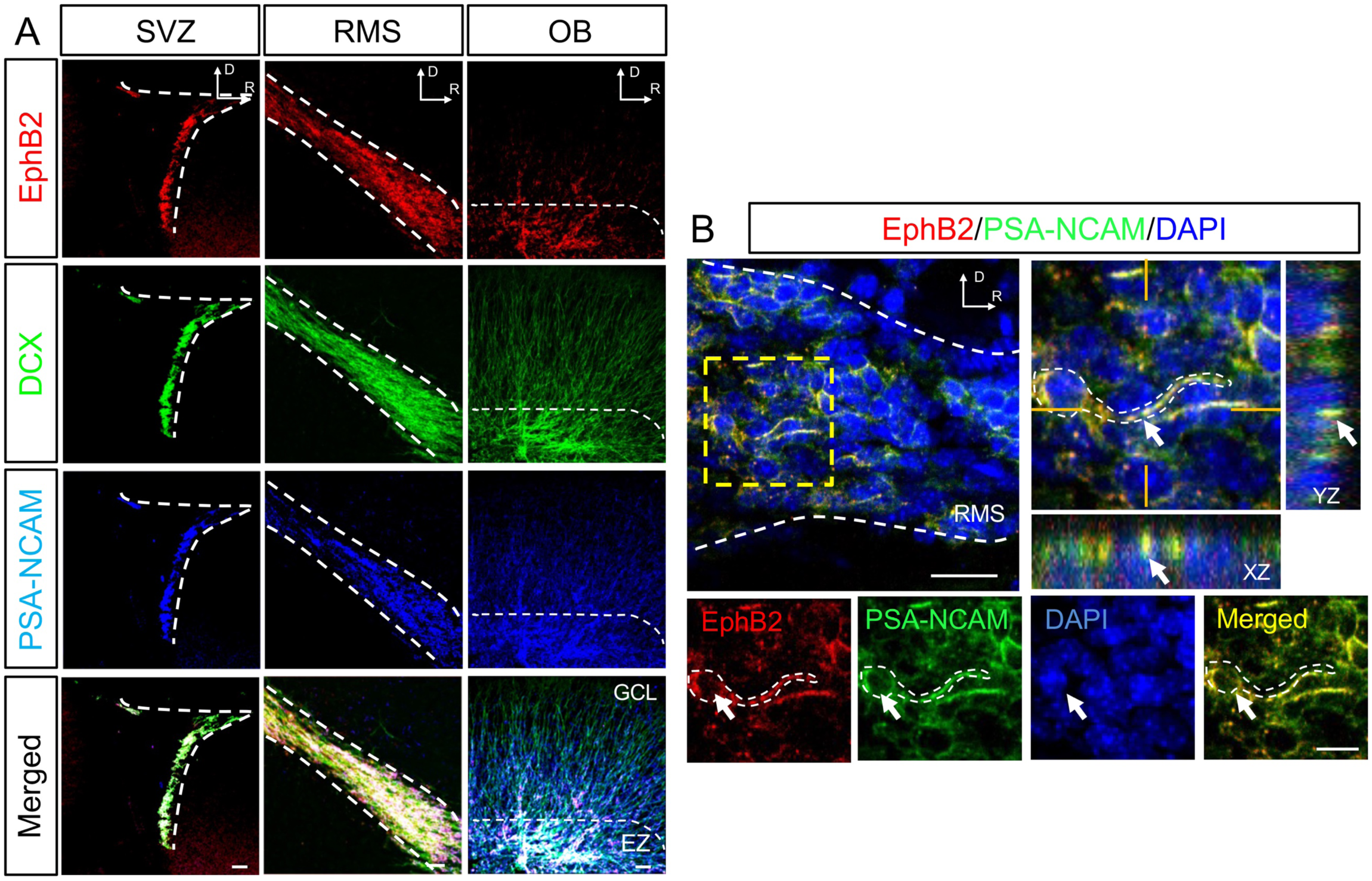
EphB2 is expressed in neuroblasts in the SVZ, RMS, and OB. (**A**) Adult neuroblasts express EphB2. Panels show α-EphB2 (red), α-DCX (green), and α-PSA-NCAM (blue). Merged images show the overlay of DCX, EphB2, and PSA-NCAM staining. From left to right: subventricular zone (SVZ), rostral migratory stream (RMS), olfactory bulb (OB). (**B**) High magnification Z-projection of EphB2 staining in the RMS. Left large panel shows merged image stained for α-EphB2 (red), α-PSA-NCAM (green), and DAPI (blue); right large image shows a magnified view of inset box (dashed lines) with projections (xz and yx) as indicated. Small panels: α-EphB2 (red), α-PSA-NCAM (green), DAPI (blue) and merged image of EphB2 and PSA-NCAM staining. Arrow indicates an example of EphB2 staining overlapping with PSA-NCAM staining in a migrating neuroblast. All orientations are labeled in the images as Dorsal (D) and Rostral (R). All images are from sagittal sections. Scale bars=50 μm in **A**, 20 μm, 10 μm in **B**.

To begin to understand the role of EphB2 in neuroblasts, we stained sections for two markers of migrating neuroblasts: DCX or PSA-NCAM. We find that EphB2 is highly expressed in migrating cells in the SVZ, the RMS, and the olfactory bulb (Fig. 1A, Supplementary Figure 2C), but not in the surrounding NeuN+ or GFAP+ cells (Supplementary Figure 3A). High-resolution imaging of migrating PSA-NCAM+ cells in the RMS confirms that EphB2 is expressed within these cells (Fig. 1B). These findings suggest that EphB2 may play specific functions in migrating neuroblasts.

EphB tyrosine kinase activation mediates cell migration and axon guidance [19, 25]. Therefore, we next asked whether EphBs are kinase active in the SVZ-RMS-OB system. Sections from wild-type (WT) mice in the SVZ-RMS-OB region were stained with our previously characterized pan phospho-EphB antibody (p*EphB) [26, 27]. Control experiments indicated that this antibody is specific for kinase active EphB in brain sections (Supplementary Figure 4). Robust p*EphB staining was found only in migrating DCX+ neuroblasts in the SVZ, RMS, and in the entry zone (EZ) of the OB (Fig. 2A). These findings indicate that EphBs are kinase active during chain migration, and may be down-regulated as cells begin radial migration in the OB.

**Figure 2.**
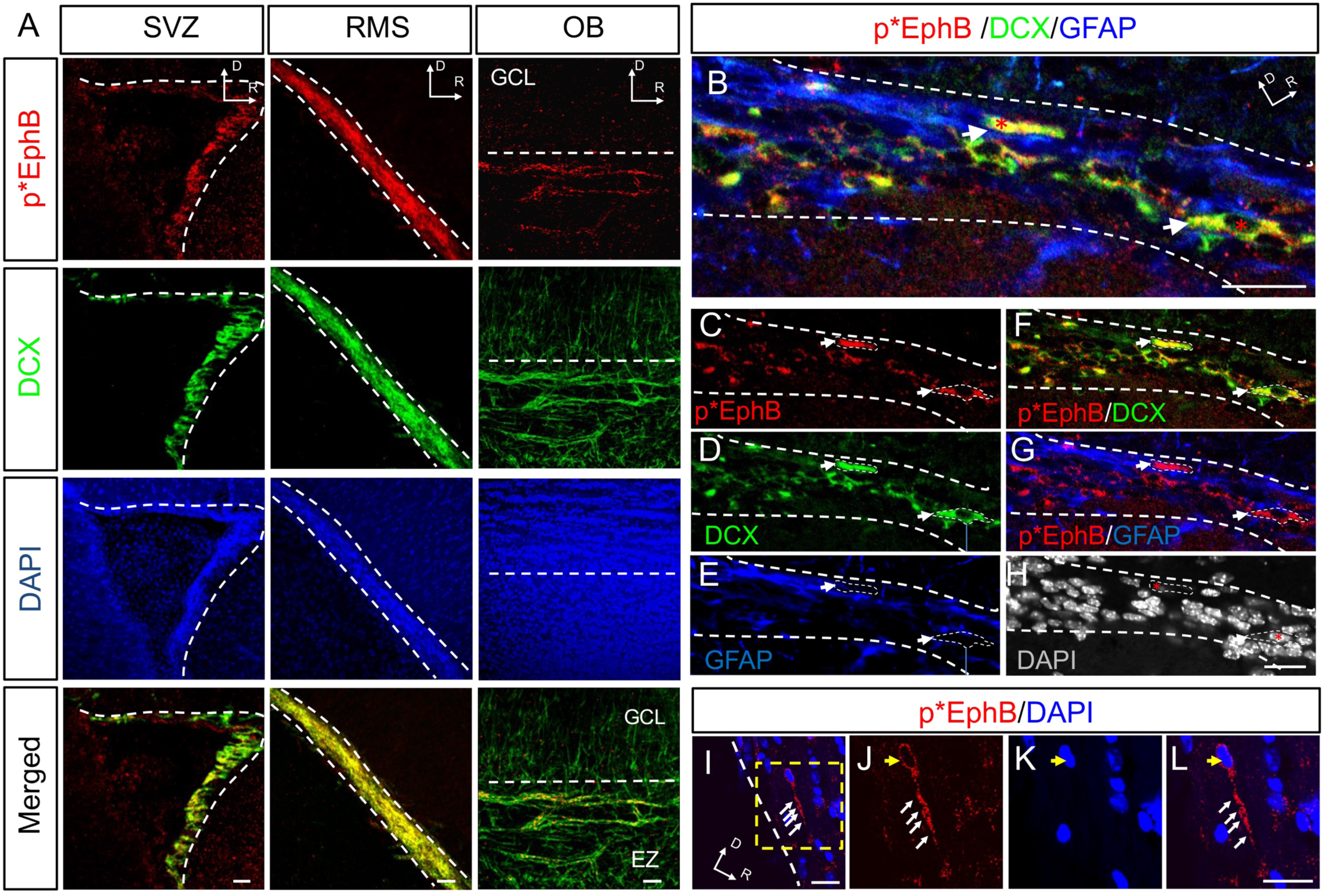
EphB2 is activated in neuroblasts in the SVZ, RMS, and entry zone of the OB. (**A**) EphB2 is phosphorylated in migrating neuroblasts. Panels show α-p*EphB (red), α-DCX (green), and DAPI (blue). Merged images show the overlay of DAPI, DCX and p*EphB staining. From left to right: subventricular zone (SVZ), rostral migratory stream (RMS), olfactory bulb (OB). The granule cell layer (GCL) and entry zone (EZ) are labeled in the OB panel and demarcated by the dashed line. (**B**-**H**) Phosphorylated EphB is found in DCX+, but not GFAP+ cells in the RMS. Dashed lines show borders of the RMS. (**B**) Merged image shows p*EphB (red), DCX (green) and GFAP (blue). (**C**-**E**) Panels show single-color images of p*EphB (red), DCX (green) and GFAP (blue) staining. (**F**-**G**) Panels show merged images of p*EphB and DCX or p*EphB and GFAP. (**H**) DAPI staining is shown in gray. Arrows in F indicate examples of DCX +/p*EphB+/GFAP-cells. (**I**-**L**) High magnification single confocal section image shows a p*EphB+(red) migrating cell in the RMS. Yellow arrow indicates DAPI+ nucleus (blue). All images are from sagittal sections. All orientations are labeled in the images as Dorsal (D) and Rostral (R). Scale bars=50 μm in **A**, 20 μm in **B**-**L**.

To examine which cells in the RMS have activated EphBs, sagittal sections containing the RMS were stained with DCX, GFAP, and p*EphB antibodies. p*EphB staining colocalized with DCX+ but not GFAP+ cells in the RMS with about 70% of migrating cells positive for p*EphB2 (Fig. 2B-H, Supplementary Figure 5). High-resolution imaging of the RMS confirms that individual neuroblasts are positive for p*EphB and indicates that p*EphB (Fig. 2 I-L). Thus, although both glia and neuroblasts in the RMS express EphB receptors, EphBs are only kinase active in the migrating neuroblasts.

To more closely examine the relationship between EphB2 expression and EphB activation as neuroblasts enter the OB and begin to differentiate, we conducted immunostaining experiments to determine the pattern of EphB2 and p*EphB in DCX expressing cells in the OB (Fig. 3A). We expect that migrating cells in the EZ of the OB will stain for both EphB2 and p*EphB, but as cells migrate into the GCL, p*EphB staining will be lost before expression of EphB2 is down-regulated. Indeed, in the entry zone of the OB before cells begin to migrate radially into the GCL, DCX+ cells stained for both EphB2 and p*EphB (Fig. 3A-E). In contrast, we rarely detected p*EphB staining in the GCL, although many cells still expressed both DCX and EphB2 (Fig. 3A and 3F-I, Supplementary Figure 6). Consistent with this model, the level of p*EphB staining remains high as neuroblasts chain migrate into the EZ. Then as cells begin to radially migrate into the GCL, the levels of p*EphB decrease rapidly (Fig. 3J-O). These findings suggest that neuroblasts maintain the expression of EphB2 until they differentiate, while kinase active EphB RTKs are found only in chain migrating neuroblasts.

**Figure 3.**
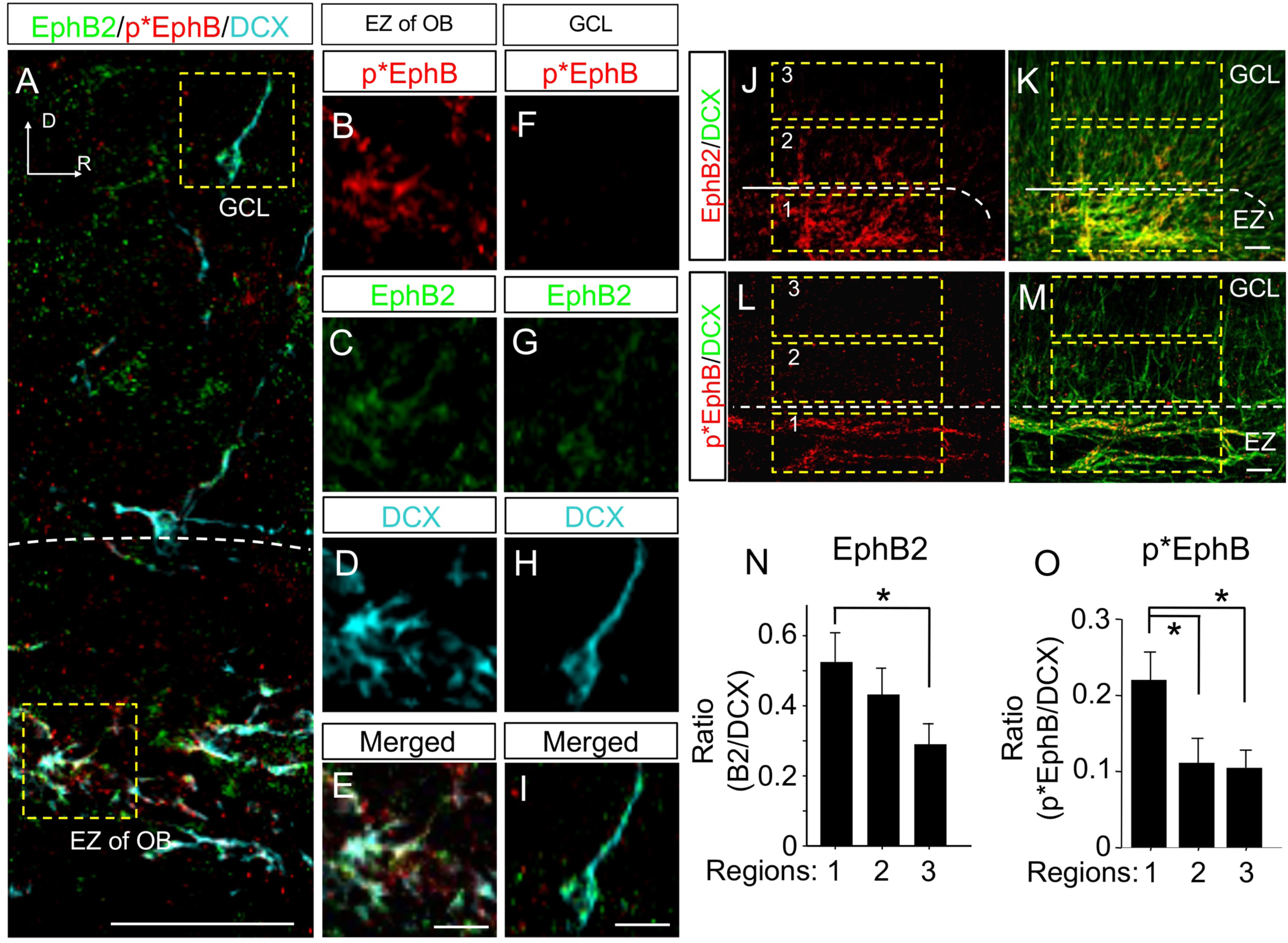
EphB kinase activity is down-regulated before EphB2 expression in the OB. (**A**) Example of EphB2 (green) and phospho-EphB (p*EphB, red) staining in cells in the entry zone (EZ) and granule cell layers (GCL) of the OB (DCX in cyan). (**B**-**E**) Magnified view of the box (yellow dashes) in the EZ showing p*EphB (red, **B**) staining for EphB2 (green, **C**) and DCX+ cells (cyan, **D**). (**E**) Merged image of **B**-**D**. (**F**-**I**) Magnified view of the box (yellow dashes) in the granule cell layer (GCL) stained as in **B**-**E**. (**J**-**K**) Example of EphB2 expression in the OB. (**J**) Shows staining for EphB2 in red. (**K**) Merged image of DCX (green) and EphB2 staining. Numbered yellow boxes show regions used for quantification shown in **N**. (**L**-**M**) Example of p*EphB expression in the OB. **L**. Image of p*EphB staining in OB (red). (**M**) Merged image of p*EphB2 and DCX (green). Boxed regions as in **J**-**K**. (**N**) Quantification of the level of EphB2 expression in different regions shown in **J**-**K** (n=5 animals). (**O**) Quantification of the level of p*EphB in regions shown in **L**-**M** (n=5 animals). Orientation is labeled in the images as Dorsal (D) and Rostral (R). All images are from sagittal sections. Scale bars=50 μm in **A** and **J**-**M**; 10 μm in bottom panels of **B**-**I**. * = p < 0.05, ANOVA.

### EphB kinase activity is essential for chain migration

EphB phosphorylation remains high in migrating neuroblasts that express EphB2 and is reduced before neuroblasts differentiate in the OB. These findings suggest that EphB kinase activity may be important in maintaining cells in a migrating, undifferentiated state. To directly determine the functional impact of EphB kinase activity, we took advantage of a Shokat mouse model: EphB1-3 triple knockin mice (TKI) in which EphB kinase activity can be selectively blocked using the synthetic inhibitor 1-NA-PP1 (Fig. 4A, [27]) without affecting EphB protein expression, uncoupling the functions of EphB protein and kinase activity. To visualize migrating neuroblasts from the SVZ, EGFP transducing lentivirus was injected into the lateral ventricle one week before interepithelial injection of 1-NA-PP1(80 mg/kg daily for one week) or vehicle control. Transduction of LV EGFP virus alone had no effects on migration of neuroblasts or differentiation of cells in the OB in WT NA-PPI injected or control injected TKI animals (Fig. 4-6). In control brains (WT control injected with 1-NA-PP1 or TKI injected with vehicle) stained for EGFP, p*EphB, and DCX, levels of p*EphB in the RMS and EGFP+/DCX+ cells was normal (Fig. 4B). In contrast, injection of 1-NA-PP1 in TKI mice resulted in a marked loss of p*EphB staining in EGFP+ cells and the RMS (Fig. 4C). These findings indicate that we can effectively block EphB kinase activity in neuroblasts *in vivo*.

**Figure 4.**
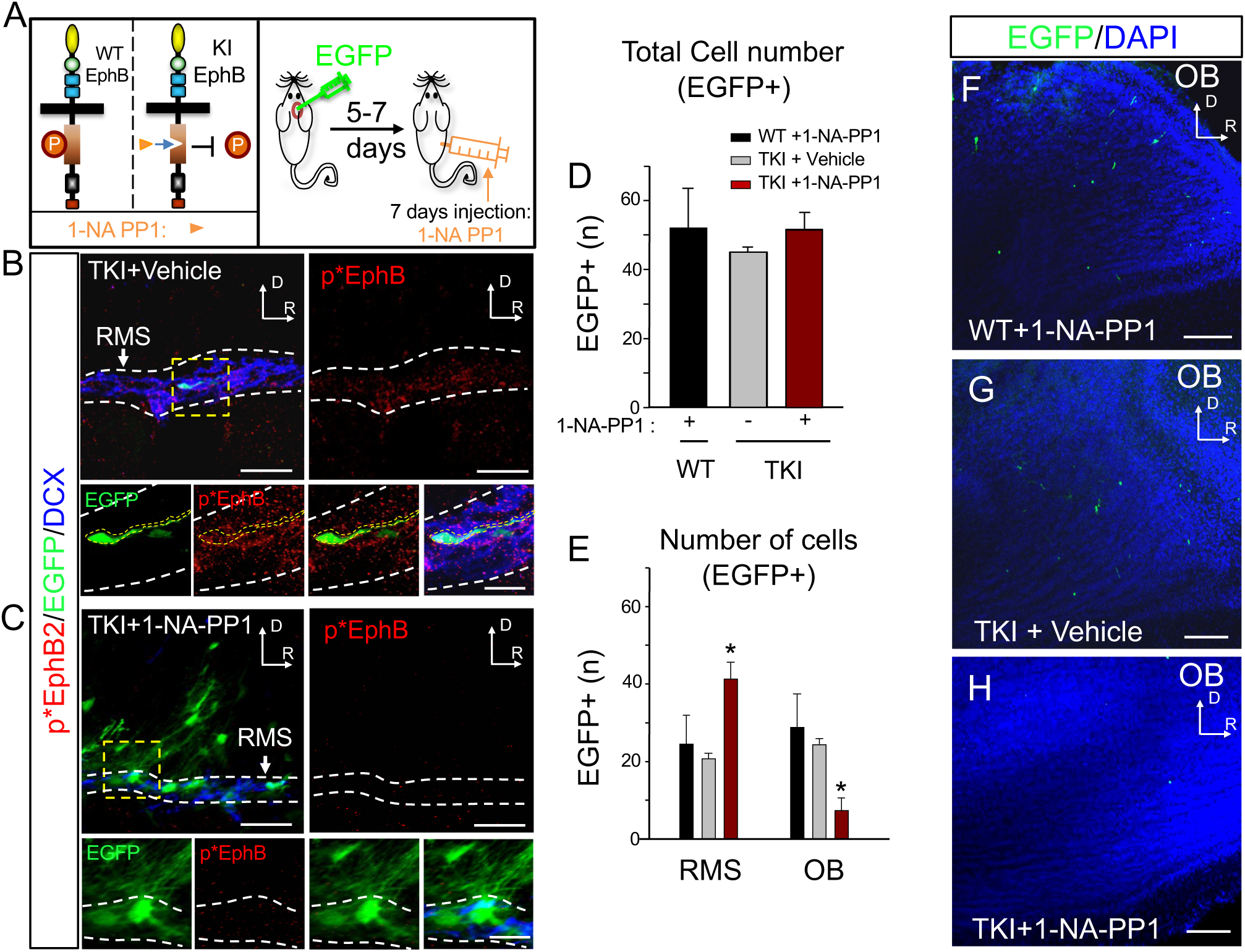
EphB kinase activity is required for migration of neuroblasts. Effects of inhibiting EphB kinase activity. (**A**) Experimental design for *in vivo* EphB kinase inhibition in TKI Shokat mice. (**B**) Example of staining for EGFP (green), p*EphB (red), and DCX (blue) in control vehicle-injected TKI animals. Large left panel shows a merged image. The large right panel shows p*EphB staining (red), and small panels show a magnified view of yellow inset box. Small panels from left to right: EGFP, p*EphB, merged panel of p*EphB/EGFP, and merged panel of DCX/p*EphB/EGFP. Labeled EGFP+ cell is outlined in yellow. (**C**) Example of staining in TKI animal injected with 1-NA-PP1 labeled as in **B**. (**D**-**E**) Quantification of migration of EGFP+ cells labeled by injection of control lentivirus into the SVZ of control or 1-NA-PP1 injected animals. (**D**) Quantification of total number of EGFP positive cells in OB and RMS (WT+1-NA-PP1: n=150; TKI+vehicle: n=145; TKI+1-NA-PP1: n=147). (**E**) Quantification of the distribution of EGFP+ cells in OB and RMS. (**F-H**) Images show EGFP+ cells in the OB of controls or 1-NA-PP1 injected animals. An injection of EGFP transducing lentivirus was made in the SVZ of animals. Two weeks later, EGFP+ cells were found throughout the SVZ, RMS, and OB. (**F**) Wild-type (WT) animals injected with 1-NA-PP1 labeled with α-EGFP (green) and DAPI (blue). (**G**) TKI animals injected with vehicle control labeled as in **F**. (**H**) TKI animals injected with 1-NA-PP1 labeled as in **I**. n=3 animals per group. All images are from sagittal sections. Scale bars=50, 20 μm in **B**-**C**; 200μm in **F**-**H**.* = p < 0.05, ANOVA.

Given the role of Eph kinases in cell and axon motility and guidance[18, 28], we hypothesized that inhibition of EphB kinase activity might impact the ability of neuroblasts to migrate. To begin to test this, we examined the number of EGFP+ cells in the RMS, and OB. The total number of EGFP+ cells in the control groups and EphB kinase blocked group were the same (WT+NA-PP1, TKI+ Vehicle, and TKI+NA-PP1, Fig. 4D). However, inhibition of EphB kinase activity resulted in a significant increase in the number of EGFP+ cells found in the RMS region and significant decrease in EGFP+ cells in the OB (Fig. 4E-I). These findings suggest that blocking EphB kinase activity results in defects in the ability of neuroblasts to migrate to the OB.

Determine the impact of inhibition of EphB kinase activity, we examined the location of EGFP+ cells in the RMS region of the brain. Surprisingly, inhibition of EphB kinase activity not only resulted in EGFP+ cells that fail to reach the OB, many EGFP+ cells were mislocalized to regions outside of the RMS (Fig. 5). To quantify the the impact of blocking EphB kinase on EGFP+ cells, we determined the position of each EGFP+ cell relative to the entry zone of the RMS (red dot) in the WT+1-NA-PP1, TKI+vehicle, and TKI+1-NA-PP1 groups (Fig. 5A). In WT+1-NA-PP1 and TKI+vehicle, nearly all virally transduced EGFP+ cells were located within the stream of chain migrating cells of the RMS (Fig. 5B-C, 5F-G), indicating that neither injection of 1-NA-PP1, nor the TKI mutation, nor transduction of SV neurons with EGFP, had any effects on the ability of neuroblasts to migrate. In contrast, in the TKI+ mice following 1-NA-PP1 injection many EGFP+ cells were located outside of the stream of chain migrating cells in the RMS and found rostral to the RMS (Fig. 5D-E, H).

**Figure 5.**
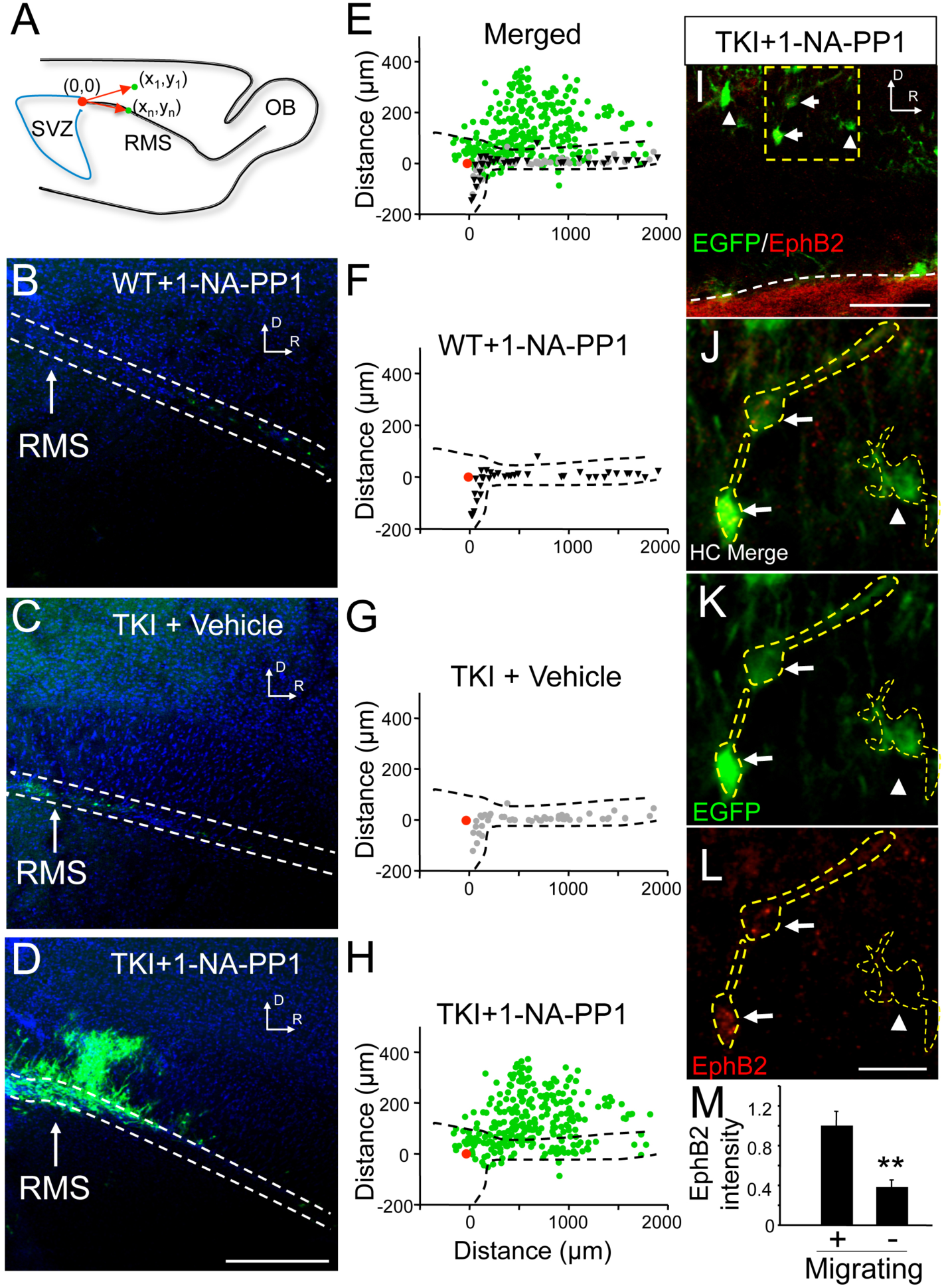
Blocking EphB kinase activity disrupts migration along the RMS. (**A**) Illustration analysis method used in **B**-**H** to determine the position of EGFP+ cells in the RMS region. (**B-H**) The pattern of EGFP+ cells labeled by injection of EGFP transducing lentivirus into the SVZ of controls or 1-NA-PP1 injected animals. (**B-D**) EGFP+ cell distribution in RMS region (arrow), labeled with EGFP (green), and DAPI (blue). (**E**) Distribution of EGFP+ cells from each of the three groups in **B-D**. Distribution of EGFP+ cells in WT+1-NA-PP1 (**F**, black, n=40), TKI+Vehicle (**G**, gray, n=48), or TKI+1-NA-PP1 (**H**, green, n=246). (**I**) TKI animals injected with 1-NA-PP1 and labeled with α-EGFP (green), α-EphB2 (red), and α-GFAP (blue). (**J**) High contrast image (HC) of merged EGFP and EphB2 staining from boxed region in **I**. (**K-L**) α-EGFP and α-EphB2 staining. Labeled EGFP+ cells are outlined in yellow. Arrows indicate EGFP+/EphB2+ cells; arrow heads indicate EGFP+/EphB2-cell. (**M**) Quantification of normalized EphB2 expression in EGFP+ cells of migrating morphology and differentiated morphology (n=30 cells/group). n=4 animals per group. Orientations are labeled in the images as Dorsal (D) and Rostral (R). All orientations are labeled in the images as Dorsal (D) and Rostral (R). All images are from sagittal sections. Scale bars = 400um in **B**-**D.**

In the TKI animals, the knockin mutations are made in all cells and approximately 70% of migrating neuroblasts are positive for p*EphB2 (Fig.2)[27]. To determine the extent of the impact of blocking EphB kinase activity on migration, we examined the size of the RMS which reflects how many cells are migrating to the OB. Consistent with the importance of EphB kinase activity in regulation of cell migration along the RMS, the both the width of the RMS measured in three sagittal sections, and the area measured in coronal sections at three different positions along the RMS, were significantly less in in TKI mice treated with 1-NA-PP1 than control (Fig. 6, Supplementary Figure 7); Sagittal sections: n=4 animals per group, normalized to width of WT: WT+1-NA-PP1: 1.0000 ± 0.0660; TKI+vehicle: 1.1237 ± 0.1414; TKI+1-NA-PP1: 0.5779 ± 0.0399; p < 0.001). These data suggest that blocking EphB kinase activity alters the ability of a large fraction of neuroblasts to migrate along the RMS, and are consistent with a model in which EphB2 kinase activity is required for chain migration and to maintain many neuroblasts within the path of the RMS.

**Figure 6.**
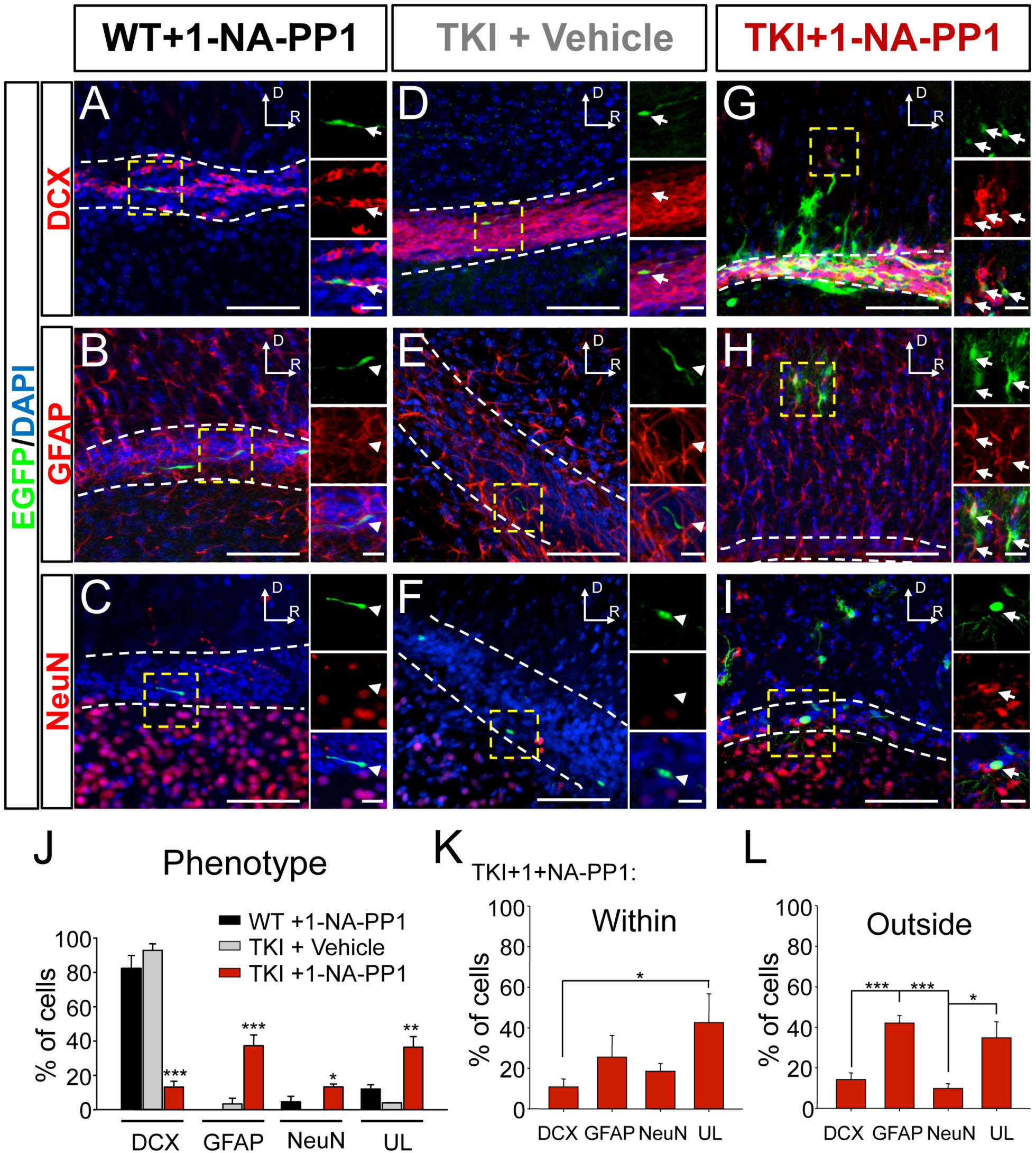
Blocking EphB kinase activity disrupts neuroblast differentiation *in vivo*. (**A**-**C**) Control experiments in WT animals labeled with different markers. (**A**) Control WT sections from mice injected with 1-NA-PP1 and labeled with α-DCX (red), α-EGFP (green), and DAPI (blue). Large panels show merged image of EGFP (green), DCX (red), and DAPI (blue). Small panels: upper right panel, EGFP; middle right panel, DCX; lower right panel, merged panel with DAPI. (**B**) Control WT sections from mice injected with 1-NA-PP1 and labeled with α-GFAP (red), α-EGFP (green), and DAPI (blue). Small panels: upper right panel, EGFP; middle right panel, GFAP; lower right panel, merged panel with DAPI. (**C**) Control WT sections from mice injected with 1-NA-PP1 and labeled with α-NeuN (red), α-EGFP (green), and DAPI (blue). Small panels: upper right panel, EGFP; middle right panel, NeuN; lower right panel, merged panel with DAPI. (**D**-**F**) Control TKI animals injected with vehicle and labeled with markers as in **A-C**. (**G**-**I)** TKI animals injected with 1-NA-PP1 and labeled as in **A-C**. (**J-L**) Quantification of the percentage of EGFP+ cells positive for DCX, GFAP, or NeuN labeled sections in WT+1-NA-PP1, TKI+Vehicle and TKI+1-NA-PP1 conditions (WT+1-NA-PP1 = 106; TKI+vehicle = 78 and TKI+1-NA-PP1 = 505; 4 animals per group). (**M**) Quantification of the proportion of EGFP+ cells positive for DCX, GFAP, or NeuN within the RMS of TKI animals injected with 1-NA-PP1 (n=220 cells). (**N**) Quantification of the proportion of EGFP+ cells positive for DCX, GFAP, or NeuN outside of the RMS in TKI+1-NA-PP1 treated animals (n=285 cells). All images are from sagittal sections. Scale bars = 100 μm, 20μm in **A**-**I.** * = p< 0.05, *** = p < 0.001, ANOVA.

We observed that EGFP+ cells with a migrating profile tended to be located nearer to the RMS, while cells with more complex morphology were located more distant from the RMS (Fig. 6 and Supplementary Figure 8). This pattern of cell morphology was similar to that seen in the OB where cells lose their migrating profile and begin to differentiate as they enter the IGL of the OB. To determine whether cells found outside of the RMS might be differentiating, we next examined the phenotype of EGFP+ cells located within the RMS or in the surrounding region. Sagittal sections were stained for EGFP and either DCX, GFAP, or NeuN. EGFP+ cells in both control groups stained only for DCX (WT+1-NA-PP1 and TKI+vehicle; Fig. 6A-6F, 6J), indicating that EphB TKI mice have normal migration in the RMS and that injection of 1-NA-PP1 alone had no impact on the phenotype of neuroblasts in the RMS. The effects of 1-NA-PP1 blockade of EphB kinase activity in the TKI mice were markedly different. In the TKI+1-NA-PP1 group, significantly fewer EGFP+ cells expressed DCX (Fig. 6G) than in the control groups (∼10% vs. ∼90%, Fig. 6J). Moreover, many EGFP+ cells expressed markers of differentiated cells (Fig. 6H-I, 6K-L). Neuroblasts that migrate through the RMS become neurons [4, 29]. Therefore, we expected that most differentiated EGFP+ cells found in the RMS would be NeuN+. However, in TKI mice following 1-NA-PP1 injections only ∼10% of EGFP+ cells were NeuN+. Surprisingly, in the TKI+1-NA-PP1 group, most EGFP+ cells found in the RMS region was GFAP+ (∼42%). These findings suggest that neuroblasts in the RMS may be less specified to the neuronal fate than previously thought.

After blockade of EphB kinase activity, many EGFP+ cells with differentiated morphology were located outside of the RMS. However, we also observed EGFP+ cells with differentiated morphologies within the RMS. To characterize the effect of blocking EphB kinase activity further, we examined the expression of cell fate markers in EGFP+ cells located both within and outside of the RMS. The fraction of DCX+/EGFP+ was similar for cells located both within and outside of the RMS (10-15% DCX; Fig. 6M-N). In contrast, a significantly higher percentage of EGFP+ cells found outside the RMS expressed a marker of mature astrocytes, with >40% expressing GFAP and only 10% expressing NeuN (Fig. 6N). Combined with the observation that blockade of EphB kinase activity led to a 50% decrease in EphB2 expression levels (Fig 5M), these findings support a model in which EphB kinase activity and expression may play distinct functions: EphB kinase activity supporting chain migration, and EphB2 expression regulating cell fate or cell differentiation.

In the OB, EphB2 kinase activity is down-regulated as cells first enter the OB and begin to differentiate, suggesting that loss of EphB kinase activity might be linked to neuroblast differentiation (Fig. 3O). To test whether blockade of EphB kinase activity might result in the premature differentiation of neuroblasts in the RMS region, we first determine the morphology of EGFP+ cells found outside of the RMS in animals where we inhibited EphB kinase activity. After blockade of EphB kinase activity (TKI+1-NA-PP1) EGFP+ cells were classified as having a migrating morphology or having a branched morphology (Supplementary Figure 8A-D). After inhibition of kinase activity, EGFP+ cells with a branched morphology, the level of EphB2 expression was significantly lower than those with a migrating profile (Supplementary Figure 8E). These findings are consistent with results from the OB and suggest a model where downregulation of EphB2 kinase activity drives radial migration and loss of EphB2 expression leads to differentiation.

### EphB2 regulates differentiation of neuroblasts

Our findings and emerging evidence indicate that the pattern of Eph and ephrin expression in the SVZ-RMS-OB is complex [14, 24]. EphB2 is expressed, and EphBs are kinase active only in migrating neuroblasts. Blocking signaling of EphBs results in defects in cell migration along the RMS. To test whether these defects are due to EphB2 expressed in migrating neuroblasts, we examined the functional significance of EphB2 in migration and cell fate determination in single cells *in vivo* using shRNA targeting EphB2 we have used and validated previously [30-32]. Injections of lentivirus encoding EGFP and either shRNA targeting EphB2 or control were made into the lateral ventricle of WT mice. After two weeks to one month, the brains of these mice were fixed, sectioned, and immunostained. Consistent with our previously published findings [30, 31], EGFP+ cells transduced with our shRNAs showed significantly reduced EphB2 expression compared to control transduced EGFP+ cells (Supplementary Fig. 10; Normalized control EphB2 expression = 1.0000 ± 0.1099, shRNA EphB2 expression = 0.1398 ± 0.0137, p < 0.01). Nestin+ stem cells found along the wall of the SVZ were infected with the injected virus and had EGFP+ progeny (Supplementary Figure 9A). Importantly, injection of either control or EphB2 shRNA transducing virus resulted in similar numbers of migrating neurons within the SVZ (Supplementary Figure 9B-C; DCX+/EGFP+ normalized to Control, Control: 1.0006 ± 0.2510 vs. EphB2 shRNA: 0.9582 ± 0.2158; p= 0.974, ANOVA; n = 4 animals in each condition). Similar to results seen when TKI mice were injected with EGFP control virus (Fig. 4), injection of control EGFP virus in WT mice had no effects on migration in the RMS or differentiation of neuroblasts in the OB (Fig. 7A-E, 8A-B,8G, 9A-B). These results indicate that we can target the cells that generate neuroblasts.

**Figure 7.**
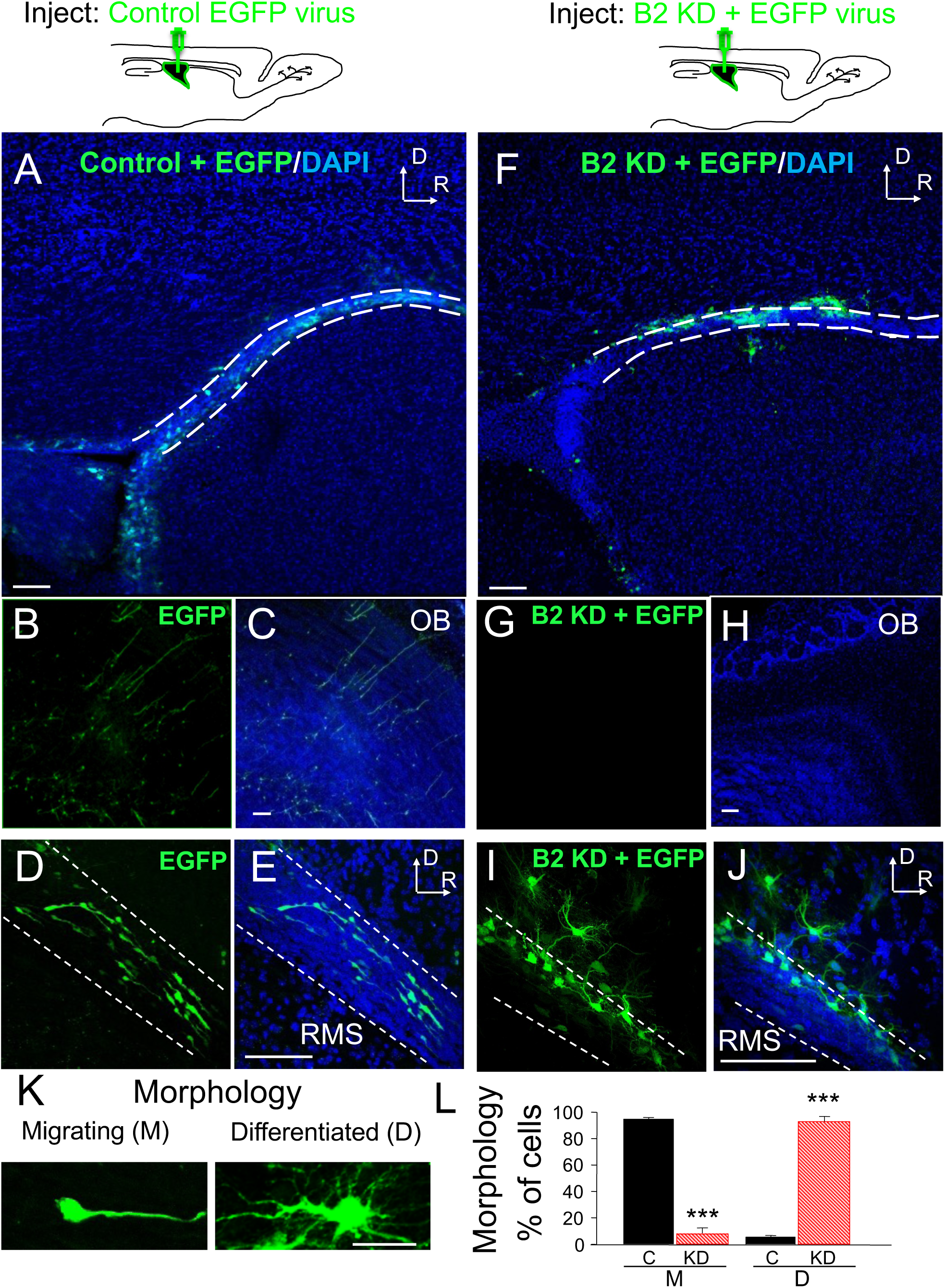
Knockdown of EphB2 disrupts migration and changes the morphology of neuroblasts. (**A-E**) EGFP+ neuroblasts are found in the RMS after injection of lentivirus transducing EGFP into the ventricle of WT mice. (**A**) Large panel stained for EGFP (green) and DAPI (blue). (**B-C**) EGFP and merged image of EGFP and DAPI staining in the OB. (**D-E**) As in **B-C** but showing RMS region. (**F-J**) EGFP+ cells in the RMS region after injection of lentivirus transducing EGFP and shRNA targeting EphB2 into the lateral ventricle of WT mice. Panels stained as in **A-E**. (**K**) Examples of the morphology of labeled cells classified as “Migrating (M)” and “Differentiated (D)”. (**L**) Quantification of the effects of control and EphB2 shRNA transducing virus on cell morphology (control n=360 cells, EphB2 knockdown n=400 cells, eight animals per group). Orientations are labeled in the images as Dorsal (D) and Rostral (R). All images are from sagittal sections. Scale bars = 100 μm in panels **A-J**, 20 μm in panel **K**, *** = p < 0.001, Student’s *t*-test.

Importantly, since relatively few cells expressed the EphB2 shRNA (Fig 7F-J), we were able to study the cell-autonomous effects of manipulating EphB2 protein expression before neuroblasts reach the OB. This approach took advantage of the stem cells present in the SVZ, which allowed labeling of cells migrating throughout the SVZ-RMS-OB system and enables examination of the effects of knocking down proteins in single cells within an otherwise normal cellular environment reducing the likelihood of compensation as seen in many knockout animals.

To determine the function of EphB2 in neuroblasts in the RMS, stereotactic injections of virus encoding either control or shRNA targeting EphB2 and EGFP were made into the lateral ventricle of eight-week-old WT mice. Based on the pattern of EphB2 expression in the OB and our results with blocking EphB kinase activity (Fig 4-6), we expected that loss of EphB2 expression would result in cessation of migration of neuroblasts. Therefore, we tested whether knockdown of EphB2 might disrupt the ability of neuroblasts to migrate into the OB (Fig 7). Neuroblasts migrate through the RMS in a few days; thus a few weeks after injection of lentivirus, in control transduced brains, many LV transduced EGFP+ neurons are found in the OB. (Fig. 7B-C). In contrast, EphB2 knockdown resulted in significantly fewer EGFP+ cells in the OB (Fig. 7G-H). These findings suggest that knockdown of EphB2 disrupts migration of neuroblasts.

EphB2 expression is down-regulated as neuroblasts undergo terminal differentiation in the OB. Therefore, if EphB2 expression is important for neuroblast differentiation, knockdown of EphB2 might result in premature stopping of neuroblasts, with cells failing to reach the OB. If this were the case, LV transduction of EphB2 shRNA might cause an increase in EGFP+ EphB2 shRNA transduced to be found either within the RMS or in the RMS region of the forebrain compared to control transduced brains. Indeed, after transduction significantly more EphB2 shRNA transduced EGFP+ cells were found within or near the RMS than in control transduced brains (Fig. 7F, 7I-J; control n = 12 animals, EphB2 shRNA n = 12 animals).

Next, to begin to examine whether downregulation of EphB2 expression might be important for differentiation, we examined the morphology of EGFP+ cells. While control LV infected EGFP+ cells had the appropriate morphology for migrating cells, only 8% of EphB2 shRNA transduced cells had a morphology consistent with a migrating cell, characterized by a long leading process followed by the soma and then a short trailing process. Instead, nearly all EphB2 shRNA transduced cells (92% of all labeled EphB2 shRNA transduced cells) had a morphology characterized by numerous branched processes and the lack of a clear leading or trailing process (Fig. 7K-L). The complex branching of EphB2 shRNA transduced cells suggests that many of these cells had begun to differentiate.

Before examining whether cells transduced with shRNA targeting EphB2 might differentiate, we examined the impact of EphB2 knockdown on the migration of cells in the RMS in more detail. To determine whether the effect of EphB2 knockdown was restricted to regions of the RMS with lower concentrations of stem cells, we labeled control and EphB2 shRNA transduced brain sections with the stem cell marker CD133. Consistent with published reports, in control brain sections CD133+ cells were concentrated within the SVZ and in the entry zone of the RMS (Fig. 8A-C). The pattern of CD133 staining was indistinguishable in the brains transduced with EphB2 shRNA (Fig. 8C and 8F). In approximately 40% of brains examined, the CD133 staining extended into the RMS, while in the remaining brains CD133+ cells remained confined within the SVZ region (Fig. 8C and 8F). Importantly, there was no correlation between the extent of CD133 labeling and lentiviral injection type.

**Fig. 8.**
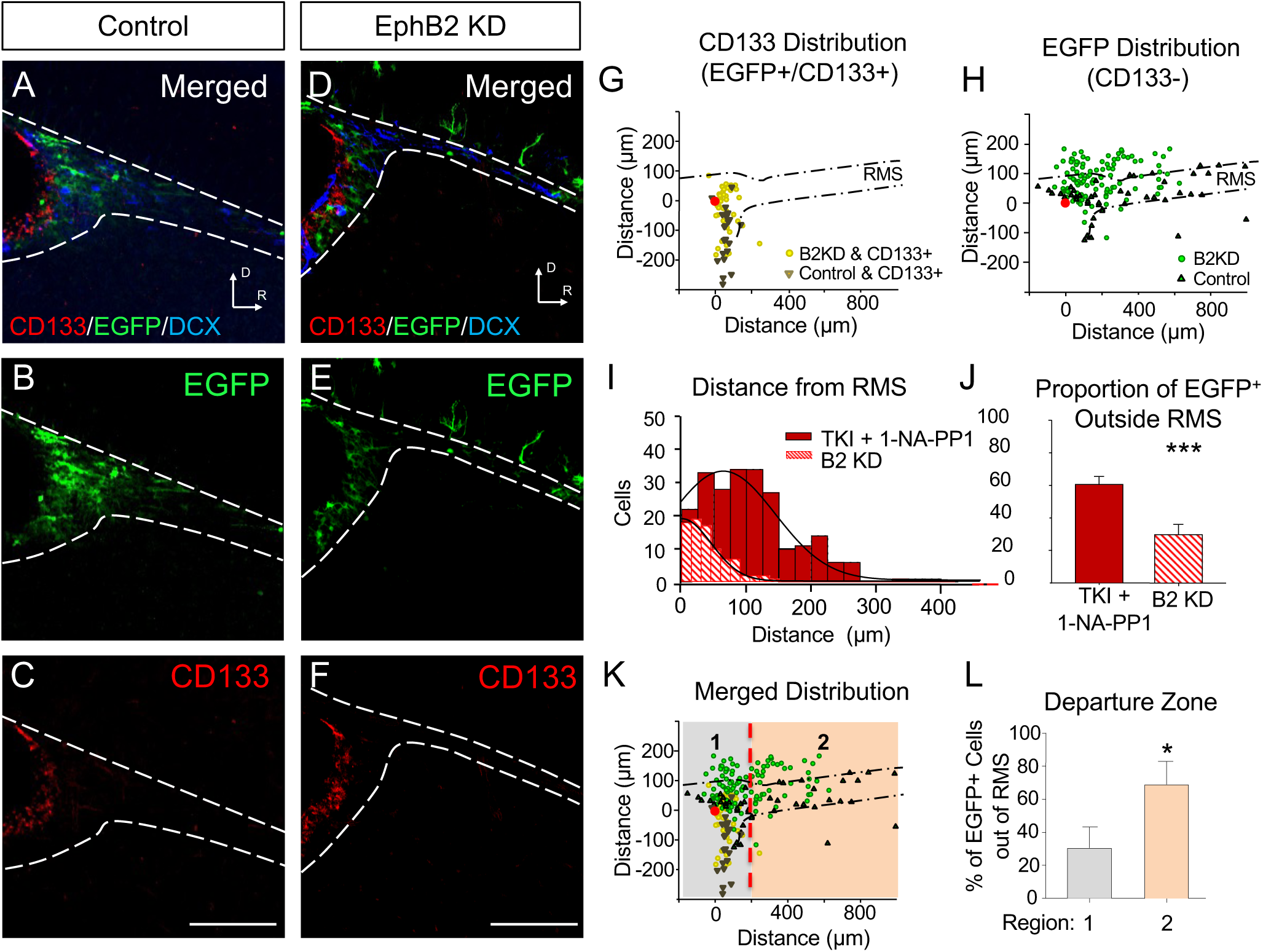
Knockdown of EphB2 expression disrupts migration of neuroblasts. (**A-C**) Control EGFP transduced brain stained for CD133 to label stem cells. (**A**) Merged image showing section stained with α-EGFP (green), α-CD133 (red), and α-DCX (blue). (**B**) EGFP (green) and (**C**) CD133 (red). (**D-F**) EphB2 knockdown (KD) transduced brain section stained to label CD133+ cells as in **A-C**. (**G**) The distribution of EGFP+ and CD133+ cells in control and EphB2 KD conditions. Location of EGFP+ cell was determined using the method illustrated in Fig. 5. (**H**) Distribution of EGFP+, CD133 negative (CD133-) cells in control and EphB2 KD conditions (B2 KD). (**I**) Histogram of the distances EGFP+ cells were located from the edge of the RMS in animals transduced with EphB2 shRNA (B2 KD; hashed bars) and after blockade of EphB2 kinase activity (TKI+1-NA-PP1; red bars; TKI+1-NA-PP1: n=227 cells, EphB2 shRNA: n= 84 cells). (**J**) Quantification of the proportion of EGFP+ cells outside RMS after TKI+1-NA-PP1 (red bar) and EphB2 shRNA (hashed bar). (**K**) The distribution of EGFP+ cells. CD133+ cells are found exclusively in SVZ region, while EGFP+ cells are found in SVZ and along RMS. The pattern of CD133 staining was used to define SVZ (**1**; gray) and RMS (**2**; peach). (**L**) Quantification fraction of EphB2 shRNA transduced cells located in the SVZ (zone 1) or RMS (zone 2) (n=206 cells; n=4 animals). All images are from sagittal sections. Scale bars = 200 μm in panels **A**-**F**, * = p < 0.05, *** = p < 0.001 Student’s *t*-test.

To determine the impact of CD133+ stem cells on the migration of cells, we measured the position of each EGFP+ cell relative to the entry zone of the RMS. EGFP+ cells were examined for co-staining with CD133. We plotted the location of each EGFP+/CD133+ cell and found that in both control and EphB2 shRNA injected groups, co-labeled cells were located only near to or within the SVZ (Fig. 8G). In contrast in the EphB2 knockdown group, EGFP+/CD133-cells were often found outside the RMS (Fig. 8H; EphB2 shRNA transduced). Thus, knockdown of EphB2 appeared not to impact the CD133+ cells but resulted in the defects in the migration of cells in the RMS.

Both blocking EphB kinase activity and knocking down EphB2 expression gave rise to EGFP+ cells found outside the RMS. Based on the pattern of EphB2 kinase activity and expression as migrating neuroblasts enter the OB, we predicted that EphB2 knockdown should result in many more EGFP+ cells being located near the RMS compared to EphB kinase blockade. To determine whether this might be the case, we measured the location of each EGFP+ cells outside of the RMS after either EphB2 knockdown or after blockade of EphB kinase activity. EGFP+ cells found outside of the RMS in TKI+1-NA-PP1 mice migrated significantly farther dorsally, on average than after EphB2 knockdown (Fig. 8I-J). These findings suggest that EphB2 expression is linked to neuroblasts migration, while EphB2 kinase activity helps to maintain chain migration of neuroblasts.

If EphB2 expression is necessary to maintain neuroblasts in a migrating state in the RMS, we would expect that knockdown of EphB2 expression should result in EGFP+ cells being found preferentially in the RMS region. To determine whether this was the case, we defined the region containing CD133+ cells as zone 1 (SVZ) and the region that did not contain CD133+ cells as zone 2 (RMS; Fig. 8K). To determine whether knockdown of EphB2 affected cells once they entered the RMS, we determined the proportion of all EGFP+ cells that exited the RMS after the CD133 border (Fig. 8K). Consistent with the model that EphB2 acts selectively within the RMS, nearly 70% of EGFP+ cells found outside of the RMS were in the zone without CD133 staining (Zone 2; Fig. 8K-L). These findings support the model that EphB2 expression is necessary to maintain neuroblast migration in the RMS.

### Loss of EphB2 drives inappropriate differentiation of neuroblasts

EphB2 expression is downregulated as cells enter the GCL of the OB, and knockdown of EphB2 expression results in neuroblasts that leave the RMS and take on mature morphology. To determine the effects of EphB2 knockdown on the cell fate in EGFP+ cells, we stained for DCX, GFAP, and NeuN and quantified the percentage of EGFP+ cells co-labeled with these markers. As expected, in control EGFP expressing brains, over 90% of EGFP+ cells in the RMS were DCX+ neuroblasts with the appropriate morphology for migrating cells (Fig. 9A and 9G). Consistent with a neuroblast phenotype, EGFP+ cells in control brains failed to stain for GFAP or NeuN (Fig. 9C, 9E, and 9H-I). In contrast, only a few EGFP+ cells transduced with EphB2 shRNA express DCX (<10%, Fig. 9B, and 9G). Instead, the majority (55%) of EGFP+ EphB2 shRNA transduced cells expressed the astrocytic marker protein GFAP (Fig. 9D, 9H, and Supplementary Fig. 11). A few (13%) cells were labeled with either NeuN (Fig. 9F, 9I, and Supplementary Fig. 11A). Thus, consistent with the pattern of EphB2 expression in the OB, after EphB2 knockdown, neuroblasts not only displayed ectopic cell morphology and disrupted migration, but they also began to express markers of mature astrocytes and neurons.

**Fig. 9.**
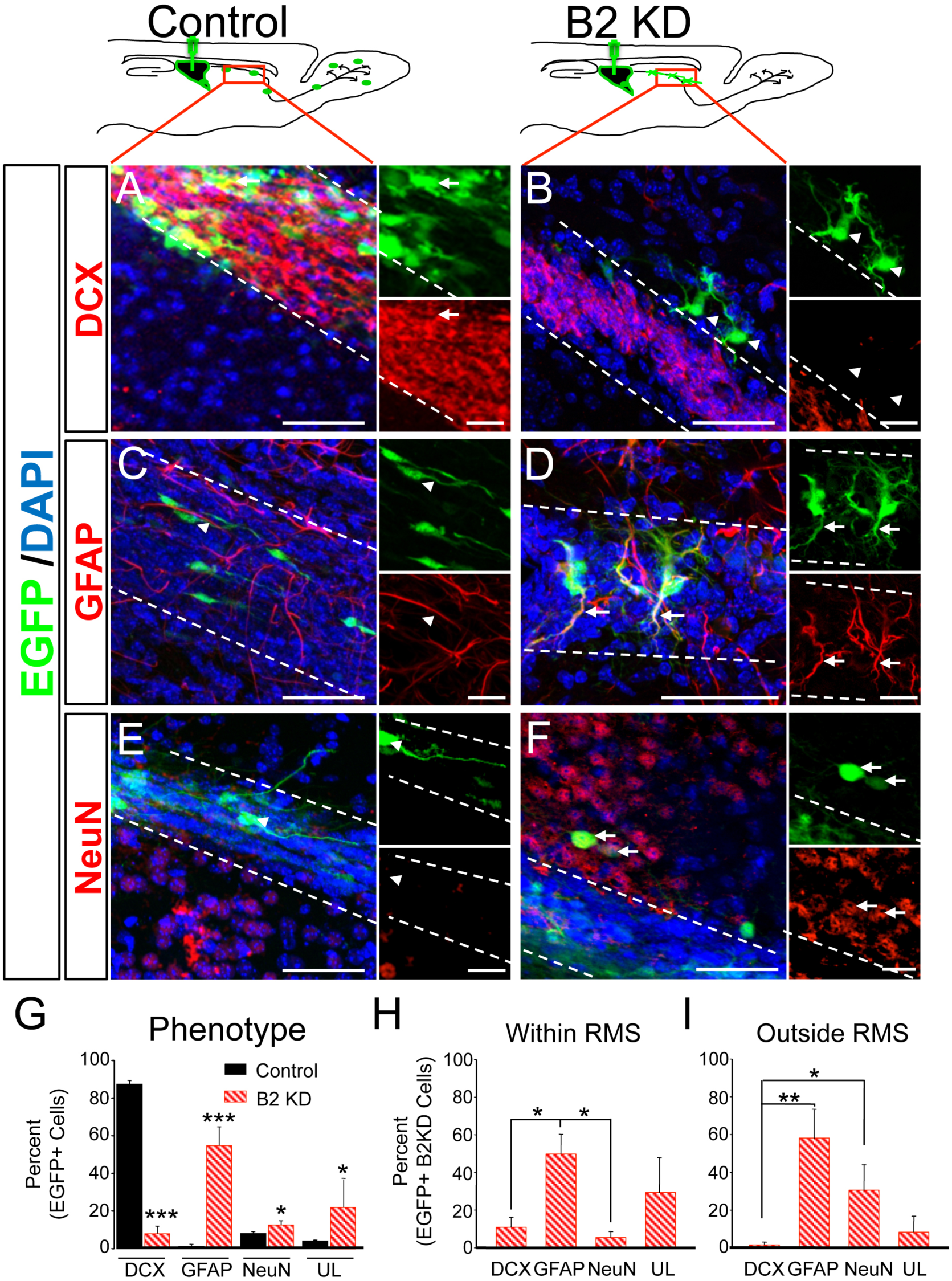
Knockdown of EphB2 promotes differentiation of neuroblasts. (**A-F**) Images of RMS stained with α-DCX (**A-B**, red), α-GFAP (**C-D**, red), and α-NeuN (**E-F**, red). Smaller panels show a magnified view of images: top right shows EGFP (green), bottom right shows α-DCX, α-GFAP or α-NeuN (red). **A**, **C**, **E** are from control injected animals. **B, D, F** are from EphB2 shRNA transduced animals (B2 KD). Few cells transduced with EphB2 shRNA were DCX+ and many cells stained for either GFAP or NeuN. Arrows indicate examples of EGFP+/DCX+, EGFP+/GFAP+ or EGFP+/NeuN+ cells. Arrow heads indicate examples of EGFP+/DCX-, EGFP+/GFAP- or EGFP+/NeuN-cells. (**G-I**) Quantification is derived from 3 different sections from 6 animals each from the control and EphB2 shRNA transduced groups. (**G-I**) Quantification of the percentage of EGFP+ cells positive for DCX, GFAP, NeuNin control and EphB2 shRNA transduced adult mouse brains (control n=690, EphB2 knockdown n=434). (**J**) Quantification of the proportion of EGFP+ cells positive for DCX, GFAP, NeuN within the RMS in EphB2 shRNA transduced mouse brains (EphB2 KD n= 286). (**K**) Quantification of the proportion of EGFP labeled cells positive for DCX, GFAP, NeuN outside the RMS in EphB2 shRNA transduced mouse brains (EphB2 KD n=148). All images are from sagittal sections. Scale bars = 50 um and 20 μm in panels **A-F**. * = p< 0.05, *** = p < 0.001, ANOVA.

In contrast to the OB, where downregulation of EphB2 expression correlates with the differentiation of neuroblasts into neurons, premature downregulation of EphB2 expression appears to cause cells to differentiate inappropriately. Since the cellular environment may contribute to cell fate, we asked whether the fate of EGFP+ EphB2 shRNA transduced cells was different depending on whether they were located within the RMS or outside of it. Consistent with a model in which knockdown of EphB2 expression results in failed migration, nearly 70% of EphB2 shRNA transduced cells remained in the RMS (Fig. 7I, 8J); of these nearly 50% were GFAP+ while less than 5% were NeuN+ (Fig. 9J). In contrast, a higher percentage of EphB2 shRNA transduced EGFP+ cells were NeuN+ when positioned outside of the RMS (∼30 vs. ∼5%; Fig. 9K), although the overall number of GFAP+/EGFP+ cells was similar to the number found in the RMS. Thus, the increased fraction of NeuN+ cells appeared to be largely due to DCX+ cells and other cells rather than to a change in the number of GFAP+ cells (Fig. 9J-K). These findings suggest that the neuronal fate of migrating cells is suppressed within the RMS. However, once cells leave the RMS and begin to move radially, a larger fraction of cells can differentiate and adopt a neuronal cell fate.

Normally, neuroblasts that migrate through the RMS and enter the OB become GAD67+ inhibitory interneurons. Therefore to further characterize the impact of EphB2 knockdown on cell fate, we stained cells for GAD67 and NeuN (Supplementary Fig. 6). Consistent with previous reports [33-35], >95% of EGFP+ cells in the OB of control brains were NeuN+ and GAD67+ (Supplementary Fig. 6A, C-G, R). In contrast, when EphB2 was knocked down only a few EGFP+ cells in the RMS region were NeuN+ and GAD67+ (<10%, Supplementary Fig 12). The majority of NeuN+ cells did not stain for GAD67 (74%). These findings suggest that even the small percentage of cells with a neuronal fate after EphB2 knockdown were either not fully differentiated or had taken an alternative neuronal fate. These data suggest that knockdown of EphB2 in single migrating cells directs the majority of neuroblasts to differentiate into GFAP+ astrocyte-like cells. Thus, before reaching the OB, migrating cells found in the RMS are likely less specified to a single fate than previously thought.

## Discussion

The molecular mechanisms controlling cell migration through the RMS have remained relatively unexplored. Here, we find that EphB2 is expressed and kinase active in chain migrating neuroblasts in the RMS and that EphB2 expression is down-regulated before neuroblasts differentiate in the OB. *In vivo* inhibition of EphB kinase activity reveals that EphB signaling regulates migration of neuroblasts along the RMS pathway to the OB while *in vivo* single cell knockdown of EphB2 indicates that EphB2 expression regulates neuroblast differentiation. Based on our findings we propose the following model. In the RMS, EphB2 expression and kinase activity maintains cells in a chain migrating state, constraining DCX+ cells to the RMS. Then as neuroblasts enter the OB, EphB2 kinase activity is lost, and cells beging to migrate radially into the OB, finally EphB2 expression is down regulated and cells differentiate. Surprisingly, our data suggest that neuroblasts in the RMS are not fully specified to a neural fate, as the knockdown of EphB2 expression and failure to migrate through the RMS, drives a majority of cells normally destined to become neurons adopt an astrocyte-like cell fate. These results suggest that, within the SVZ-RMS-OB migratory pathway, cell-cell interaction either between neuroblasts and/or the ensheathing glial cells are integral to establishing a neural identity.

Interactions between neuroblasts and the ensheathing glial cells have long been thought to enable the precise migration of cells from the SVZ to the OB [1-3, 16]. However, the study of these glial cells has focused on their potential role as a mechanical barrier to migrating cells [11, 12]. Our data indicate that EphB2 is kinase active within migrating DCX+ cells. EphB2 activity is downregulated in DCX+ cells as they enter the OB and begin migrating radially. Consistent with these data, artificially down-regulating EphB kinase activity in migrating neuroblasts results in a premature radially migration cells from the RMS. These data support a model in which radial migration is triggered by the loss of EphB2 phosphorylation in neuroblasts as they enter the OB.

In the SVZ, EphB and ephrin-B were implicated in migration by infusing the extracellular domains of EphBs or ephrin-Bs into the lateral ventricle (Conover et al., 2000). Infusion of EphB2-FC or ephrin-B2-FC, but not ephrin-B1-FC, disrupted chain migration within the SVZ. Although not appreciated at the time, the finding that ephrin-B2 but not ephrin-B1 causes defects in migration suggests that the EphB protein responsible for the migration defects in the SVZ is not EphB1, EphB2, or EphB3. EphB1-3 bind ephrin-B1 and ephrin-B2 with similar affinities, therefore one would expect that infusion of these ligands would give similar phenotypes. EphA4 also binds both ephrin-B1 and ephrin-B2 with similar affinities [36-38]. In contrast, EphB4 binds ephrin-B2 preferentially [39, 40]. EphB4 is an essential gene expressed principally in the vasculature [41], and little studied in the CNS. Although there is little evidence supporting high levels of EphB4 expression in the SVZ, it will be interesting to determine whether EphB4 is expressed in the SVZ at low levels to regulates migration of cells. Regardless, our data suggest that EphB2 is responsible for migration in the RMS, suggesting that different EphBs may regulate migration of neuroblasts in specific regions of the SVZ-RMS-OB system.

For migrating cells to reach their targets, they must move at an appropriate rate and follow the correct route. Several lines of evidence suggest that EphBs are likely to be important for cell migration: EphBs have a demonstrated role in migration of intestinal progenitors, neural crest cells, the dental pulp, and hippocampal neuroblasts [20, 42-45]; and *in vitro* ephrin-B1/EphB2 interactions promote migration in HEK 293T cells [46]. We find that EphB kinase activity is linked to the ability of migrating neurons to follow the correct route along the RMS to the OB. In the SVZ-RMS-OB system, Slit/Robo, PSA-NCAM, VEGF, and the neuregulins appear to be essential for enabling neuroblasts to move at the proper rate [15, 16, 47-50].

Manipulation of the Slit/obo, PSA-NCAM, VEGF, and the neuregulin signaling systems reduce the rate of migration, as evidenced by increased numbers of migrating cells found within RMS. However, such manipulation does not appear to alter the ability of cells to follow the correct pathway, cause them to stop migration, or alter the fate of the cells. These earlier findings suggested that the glial cells that line the RMS might form a physical barrier. Our data challenge this hypothesis since single-cell *in vivo* knockdown of EphB2 in neuroblasts and blockade of EphB signaling causes profound migration defects and results in fewer cells that can follow the correct path along the RMS. Thus, the glial sheath surrounding the RMS only poses a barrier to migrating cells if the EphB kinase can be activated. Importantly, EphB2 knockdown does not appear to affect migration within the SVZ, suggesting EphB2 may function selectively in the RMS.

Our data suggest that the kinase activity of EphB may maintain cells in the chain migrating state or mode. Reelin appears to regulate detachment of neuroblasts in the OB [51, 52], and recent findings indicate that EphB2 binding to Reelin down-regulates EphB and ephrin-B signaling [53-55]. It is interesting to consider that the EphB-Reelin interaction might be responsible for the changes that we observed in EphB2 kinase activity as neuroblasts enter the OB and begin to migrate radially. Consistent with this model, EphB2 is expressed as neuroblasts migrate into the GCL, but EphB2 kinase activity is down-regulated as cells migrate radially to leave the RMS and enter the zone where Reelin is expressed. Migration appears to stop, and cells begin to differentiate once EphB2 expression is lost. An important question will be to determine the source or sources of ephrin ligands within the RMS and OB. Indeed, Ephs and ephrins appear to have a complex pattern of expression in the RMS, with data suggesting that several EphB and ephrin ligands are expressed in the ensheathing glial cells [14, 24]. It is tempting to speculate that repulsive interactions between glial-expressed ephrin and neuroblast-expressed Eph might explain both the path followed by neuroblasts and the high level of EphB2 kinase activity seen in migrating cells. Regardless, our findings suggest that EphB kinase signaling, rather than simple expression levels, may regulate neuroblast migration, while downregulation of EphB2 expression may trigger neuroblast differentiation.

Unexpectedly, knockdown of EphB2 expression results in the premature and inappropriate differentiation of neuroblasts. It is well known that cells generated in the SVZ are destined to become neurons when they reach the OB. Indeed, deletion of either PTEN or Tsc1 results in cells that fail to migrate to the OB but still differentiate into neurons [56, 57]. However, after EphB2 knockdown, the majority of neuroblasts fail to migrate properly and change fate to become GFAP+ cells. These findings indicate that neuroblasts born in the SVZ may be less committed to a specific neural fate than previously appreciated [29, 58], that migration through the RMS is linked to the cell fate of OB interneurons, and that EphB2 regulates neuronal cell fate. Importantly results from both knockdown of EphB2 and blockade of EphB kinase activity generate similar phenotypes. Interestingly, downregulation of EphB2 expression in the colon results in gene expression that drives cell differentiation [59], suggesting that EphBs may be important regulators of cell maturation or fate. However, more work will be needed to determine the specific role EphB2 plays as neuroblasts enter the OB.

In the absence of EphB2, cells stop (70% inside the RMS) and differentiate into GFAP+ and NeuN+ cells. In contrast, if EphB kinase activity is only blocked, cells appear to keep migrating but do so inappropriately. Indeed, after a kinase blockade, 60% of cells fail to respect the RMS boundary and migrate radially. Blockade of EphB activity results in cells that migrate further than cells expressing EphB2 shRNA (Fig. 8I). Similarly, phosphorylation of EphB in the OB is down-regulated as neuroblasts migrate radially, but EphB2 expression is lost as the cells differentiate. We propose that EphB2 kinase activity is required to maintain chain migration through the RMS. In the absence of EphB kinase activity, neuroblasts migrate radially and differentiate once EphB2 expression is down-regulated.

Our findings suggest that migration through the RMS is likely to be mediated by a complex set of regulatory and guidance cues, not physical constraints, which may also have functions in determining cell fate. By defining a single molecular mechanism that both constrains cells along a specific migratory pathway and regulates the differentiation of these cells, our data show, unexpectedly, that migration is coupled to the process of cell fate determination.

## Materials and Methods

### Animals and Stereotactic Injections

All animal procedures were performed in strict accordance with the NIH Guide for the Care and Use of Laboratory Animals and were approved by our Institutional Animal Care and Use Committee. Two month old WT CD1 (Charles River), EphB1-/-, 2-/-, 3-/- mice or EphB TKI mice were anesthetized with 0.03 mL of a mixture of ketamine (90.9 mg/mL) and xylazine (9.1 mg/mL). In all cases virus was diluted in PBS to 1E+10 IFU/mL and 1μL was delivered at a rate of 200 nL/min using a programmable syringe pump (World Precision Instruments). For all controls U6.empty.CMV.EGFP was used. All viruses were generated at the Penn Vector Core. For EphB2 Knockdown: either control or U6.ShmEphB2.CMV.EGFP (EphB2 knockdown virus) was injected at coordinates targeting the lateral ventricle: A/P: 0.0 mm, M/L: +0.8 mm, and D/V: 2.0 mm. For EphB TKI mice: EGFP expressing virus was injected at coordinates targeting the lateral ventricle: A/P: 0.0 mm, M/L: +0.8 mm, and D/V: 2.0 mm.

### Delivery of 1-NA-PP1

Experiments using the EphB TKI mice were conducted using methods similar to those described previously [27]. Briefly, 5-7 days after virus injection, WT mice and EphB TKI mice were injected with either vehicle (10% DMSO, 20% Cremaphor-EL, and 70% saline) or 1-NA-PP1 (80mg/Kg, Cayman Chemical Company) every 12 hours for 7 days. Mice were injected either subcutaneously or intraperitoneally, with both methods giving similar results.

### Immunostaining and Microscopy

For immunostaining of brain sections, mice were transcardially perfused with ice-cold PBS followed by 4% paraformaldehyde in PBS. Brains were removed, fixed overnight in 4% paraformaldehyde at 4°C and incubated in 30% sucrose at 4°C until equilibrated. Brains were frozen for 1 min at −50°C in dry ice cooled 2-methylbutane (Fisher Scientific) and stored at −80°C until sectioned. 40 μm coronal or sagittal sections were cut using a freezing sliding microtome and processed as free-floating sections. Primary and secondary antibodies were incubated in PBS with 0.4% TX-100 and 5% normal goat or donkey serum for 24 h at 4°C. Primary antibodies were the following: mouse α-GFAP (1:500, Sigma 3892), mouse α-NeuN (1:250, Chemicon MAB377), rabbit α-doublecortin (1:300, Cell Signaling 4604), rabbit α-GFAP (1:500, Sigma 9269), goat α-EphB1, goat α-EphB2, goat α-EphB3 (1:100-200, R&D Systems), rat α-CD133 (1:200, Millipore MAB 4310), mouse α-GAD67 (1:1000, MAB5406). Secondary antibodies were the following: goat α-rabbit 546 (1:250, Invitrogen), goat α-rabbit 633 (1:250, Invitrogen), goat α-mouse 546 (1:250, Invitrogen), goat α-mouse 488 (1:250, Invitrogen), goat α-rat 546 (1:250, Molecular Probes), donkey α-rabbit Cy3 (1:250, Jackson Lab), donkey α-rabbit 633 (1:250, Jackson Lab), donkey α-mouse Cy3 (1:250, Jackson Lab), donkey α-mouse Cy3 (1:250, Jackson Lab), donkey α-rat Cy3 (1:250, Jackson Lab), donkey α-goat Cy3 (1:250, Invitrogen). Secondary antibodies were applied for 2 h at room temperature and sections were then washed for 10 min with 1x PBS 3 times. Sections were mounted on pre-coated slides (Superfrost Plus Microscope Slides, Fisher) and coverslipped using a DNA counterstain (4’,6-diamidino-2-phenylindole; DAPI, Vector Labs) to label nuclei.

### Validation of Phospho-specific anti-EphB antibody

Phospho-specific α-EphB antibody against the partial peptide sequence of the EphB2 intracellular region [26] was used for detecting activated EphB2 in brain sections. Preabsorption control for this antibody was performed using a phosphorylated intracellular protein fragment of EphB2. Autophosphorylation of the intracellular fragment of EphB2 (N-terminal His_6_-tagged recombinant human EphB2 residues 560-end, EMD Millipore, Billerica, MA) was validated by incubating 3.1 μg EphB2 intracellular protein fragment in a solution of 8 mM MOPS/NaOH (pH 7.0), 0.2 mM EDTA, 10 mM MnCl_2_, 100 μM ATP, and 10 mM MgAc at 30°C for 30 min. The efficacy of phosphorylation was confirmed by western blotting using a part of the reaction product (Supplementary Fig. 3). Indicated primary antibodies were presented in blocking solution for 2 h at room temperature or overnight at 4°C: rabbit polyclonal α-phospho-EphB (1:1,000, [26]), mouse monoclonal α-EphB2 (1:500, Invitrogen). For preabsorption, the phosphorylated EphB2 fragment was added to the working solution of phospho-specific α-EphB antibody at a concentration of 4 μg/ml at 4°C overnight. For detecting active EphB2 in brain slices, antibody solution was preincubated with EphB2 intracellular protein fragment in the same solution without ATP and MgAc was used to reduce the signals that are not specific for phosphorylated status.

### Quantification of Cell Types

To quantify the number of cells, 0.09-0.5 μm step confocal z-series of brain sections were analyzed using MetaMorph® image analysis software (Molecular Devices). Cells identified by DAPI staining were determined to be either DCX+, GFAP+, NeuN+, or unlabeled. Additionally, the morphology of each cell was determined to be migrating or differentiated. The boundaries of the RMS were determined based on the cellular density as visualized by DAPI staining and cells were determined to be either within or outside of the RMS. Cells at the edge of the field of view were excluded from analysis.

### SVZ Whole-Mount Assay

Two month old CD-1 mice were injected with EGFP virus and SVZ whole-mounts of striatal lateral wall were dissected 2 weeks after injection and processed as described previously [60]. Briefly, the SVZ whole-mounts were fixed overnight in 4% paraformaldehyde in 0.1M PBS with 0.1% T-X100 at 4°C, washed with PBS, and blocked with 10% normal goat or donkey serum in 0.1 M PBS with 0.1% Triton-X100 for 1 h at 4°C. Whole-mounts were then incubated with primary antibodies diluted in blocking buffer for 24-48h at 4°C. Initially, primary antibodies are washed off by 2 quick rinses in PBS with or without 0.1% Triton-X100. Slides were then washed 3 times for 20 min each at room temperature and incubated with appropriate secondary antibodies at 4°C for 24-48 h. After incubation with secondary antibody, slides were washed again, as above, and coverslipped with aquamount (PolySciences).

### Measurement for the width of RMS

Doublecortin (DCX) is used as marker for the width measurement of RMS. The same number of mice (n=4 animals per group) was used for the WT+1-NA-PP1, TKI+vehicle and TKI+ 1-NA-PP1 groups. All sections are Z-merged by the same setting, and from the same sites of the RMS (beginning, middle and end parts). The number of sections (n=3) is same in each part for all 3 groups (total n=12 per group). Each width measurement is normalized to the mean of the WT+1-NA-PP1 group.

### Statistical Analyses

All data were expressed as means ± SEM and analyzed using ANOVA and Student’s *t* test. The level of significance for all analyses is as noted in the figure legends or main text. Distribution of the data was assumed to be normal, but this was not formally tested. No statistical methods were used to predetermine sample sizes, but our sample sizes are similar to those reported in previous publications [26]. Probability values of less than 5% were considered statistically significant. Unless stated otherwise, statistical measures were conducted on a per-cell basis, collected from a minimum of three independent experiments.

| REAGENT or RESOURCE | SOURCE | IDENTIFIER |
| --- | --- | --- |
| **Antibodies** |  |  |
| Mouse anti-GFAP | Sigma-Aldrich | Cat# G3893; RRID: AB_477010 |
| Rabbit anti-GFAP | Sigma-Aldrich | Cat# G9269; RRID: AB_477035 |
| Mouse anti-NeuN | Millipore | Cat# MAB377; RRID: AB_2298772 |
| Rabbit anti-Doublecortin | Cell Signaling Technology | Cat# 4604; RRID: AB_10570650 |
| Goat anti-EphB1 | R&D Systems | Cat# AF467; RRID: AB_355262 |
| Goat anti-EphB2 | R&D Systems | Cat# AF467; RRID: AB_355262 |
| Goat anti-EphB3 | R&D Systems | Cat# AF4467; RRID: AB_2107593 |
| Rat anti-CD133 | Millipore | Cat# MAB4310; RRID: AB_11212378 |
| Mouse anti-GAD67 | Millipore | Cat# MAB5406; RRID: AB_2278725 |
| **Chemicals, Peptides, and Recombinant Proteins** |  |  |
| Ketamine | Sigma-Aldrich | CAS: 6740-88-1 |
| Xylazine | Sigma-Aldrich | CAS: 7361-61-7 |
| 1-NA-PP1 | Cayman Chemical Company | Cat# 14320 |
| Cremophor EL | Sigma-Aldrich | Cat# C5135 |
| Recombinant Human EphB2 Fragment | EMD Millipore | Cat# 3295 |
| Biological Samples |  |  |
| Brain tissues from mice | This paper | N/A |
| **Experimental Models: Organisms/Strains** |  |  |
| Mouse: CD1 (WT) | Charles River Laboratories | RRID: IMSR_CRL:22 |
| Mouse: EphB1<sup>-/-</sup> | Previous study | [Reference if available] |
| Mouse: EphB2<sup>-/-</sup> | Previous study | [Reference if available] |
| Mouse: EphB3<sup>-/-</sup> | Previous study | [Reference if available] |
| Mouse: EphB TKI | Generated as per Soskis et al., 2012 | [Reference if available] |
| **Recombinant DNA** |  |  |
| U6.empty.CMV.EGFP virus | Penn Vector Core | N/A |
| U6.ShmEphB2.CMV.EGFP virus | Penn Vector Core | N/A |
| **Software and Algorithms** |  |  |
| MetaMorph Imaging Software | Molecular Devices | RRID: SCR_002368 |
| NIH ImageJ | NIH open source |  |
| **Other** |  |  |
| Programmable Syringe Pump | World Precision Instruments | Cat# UMP3 |
| Superfrost Plus Microscope Slides | Fisher Scientific | Cat# 12-550-15 |
| VECTASHIELD Mounting Medium with DAPI | Vector Laboratories | Cat# H-1200; RRID: SCR_001881 |

## Author Contributions

W.Z. and J.M. conducted the majority of the experiments shown and undertook data analysis. K.H. conducted experiments related to the EphB2 phospho-specific antibody. H.H. provided advice on the use of the TKI mice. All authors discussed the results and commented on the manuscript. M.B.D, W.Z., and J.M. designed the experiments and wrote the manuscript.

## Acknowledgments

We thank Rachel Hodge and the other members of the Dalva laboratory for helpful discussions and advice; Dr. Michael Greenberg (Harvard) for generously supplying the EphB TKI mice and phospho-EphB antibody; Dr. Douglas Coulter (U. Penn) for use of his confocal microscope; Dr. Mark Henkemeyer (UTSW) for the EphB TKO mice. The Neural Developmental Disabilities Training Grant (CHOP, T32 NS007413) and Alavi-Dabiri Postdoctoral Fellowship supported J.M. NIH grants from NIDA (DA022727), and the NIMH (MH086425 and MH100093) and a grant from the Dana Foundation to Dr. Dalva supported this work.

## Ethics

All animal procedures were performed in strict accordance with the NIH Guide for the Care and Use of Laboratory Animals and were approved by the appropriate Institutional Animal Care and Use Committee.

