## Supplemental Figures and Legends for "Specific neuroblast-derived signals control both cell migration and fate in the rostral migratory stream"

**Supplementary Figures**

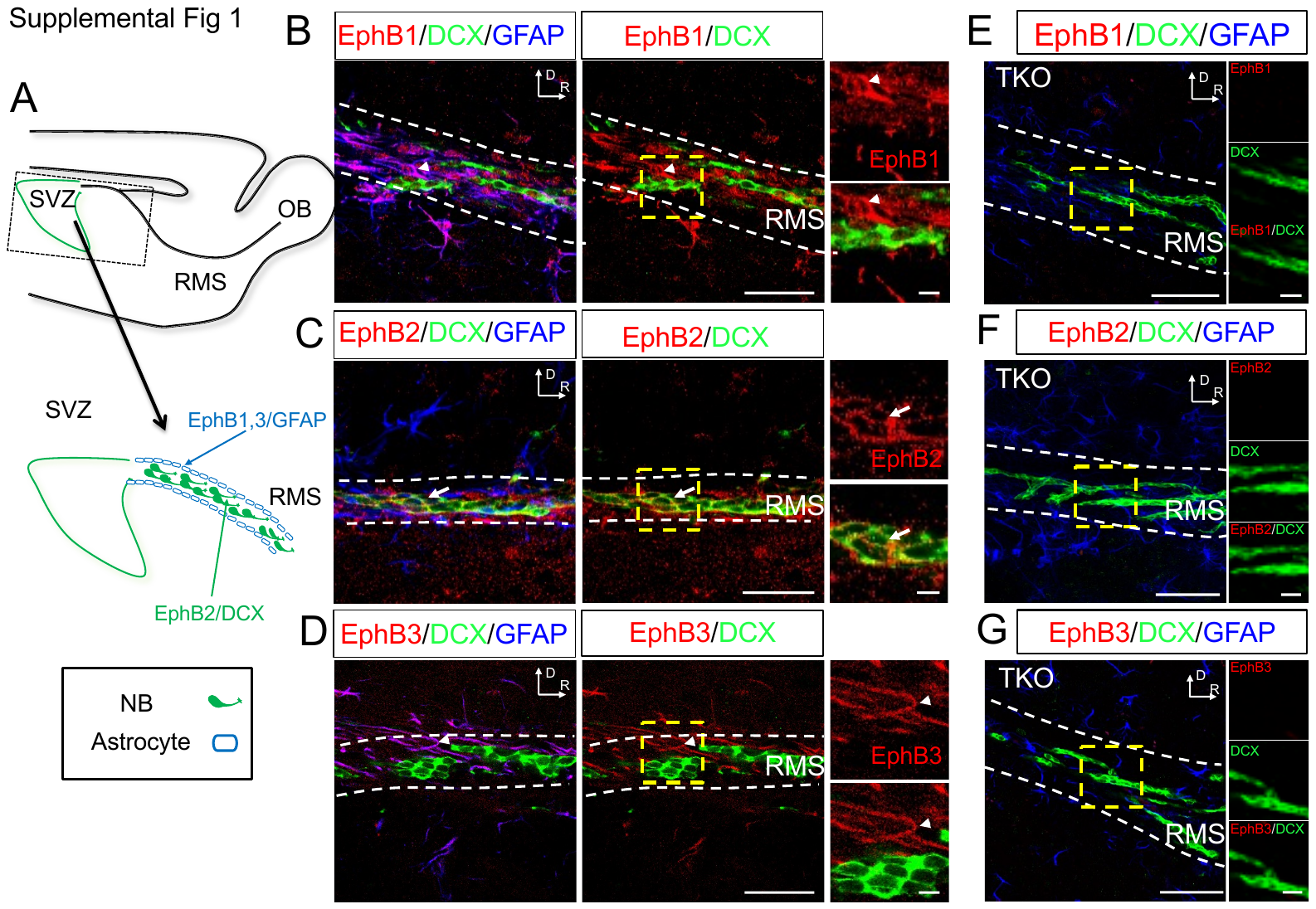

**Supplementary** F**igure** 1. **Pattern of expression of EphB1, EphB2, and EphB3 in the RMS and ensheathing astrocytes.** (**A**) Model illustrating the pattern of expression of EphB1, EphB2 and EphB3 in the RMS region of the rodent brain. EphB2 expression is found in DCX positive cells (green) while EphB1 and EphB3 are expressed in GFAP positive cells (blue). (**B**) A WT mouse brain section stained with α-EphB1 (red), α-DCX (green) and α-GFAP (blue). Left large panels shows merged image of EphB1 (red), DCX (green) and GFAP (blue) in RMS. Middle large panels shows merged image of EphB1 (red) and DCX (green). Right small images show magnified view of inset box (dashed lines): EphB1 (red) and a merged image of EphB1 and DCX (red+green). (C) Images stained as in B, but with α-EphB2. (D) Images stained as in B, but with α-EphB3. Arrow (C) indicate an example of EphB2+/DCX+ cell, and arrow heads indicate examples of EphB1+/DCX- (B) and EphB3+/DCX- cells (D). (**E**-**G**) Immunostaining in the EphB1-3 triple knockout mouse (TKO) validating antibody specificity. EphB1-3 triple knockout mice (EphB TKO; EphB1^-/-^, EphB2^-/-^, EphB3^-/-^) mouse brain sections stained with EphB antibodies used in **B**-**D**. Immunostaining was conducted in parallel with experiments in WT mouse brains and the same confocal setting were used. (**E**) Large panel shows merged image of α-EphB1 (red), α-DCX (green), and DAPI (blue). Right small images show magnified view of inset box (dashed lines): α-EphB1 (red), α-DCX (green) and merged image of EphB1 and DCX (red+green). (**F**) As in **E**, but stained with α-EphB2. (**G**) As in **E**, but stained with α-EphB3. Orientations are labeled in image as Dorsal (D) and Rostral (R). All images are from sagittal sections.  Scal bars=100 μm, 20 μm in **B**-**G.**

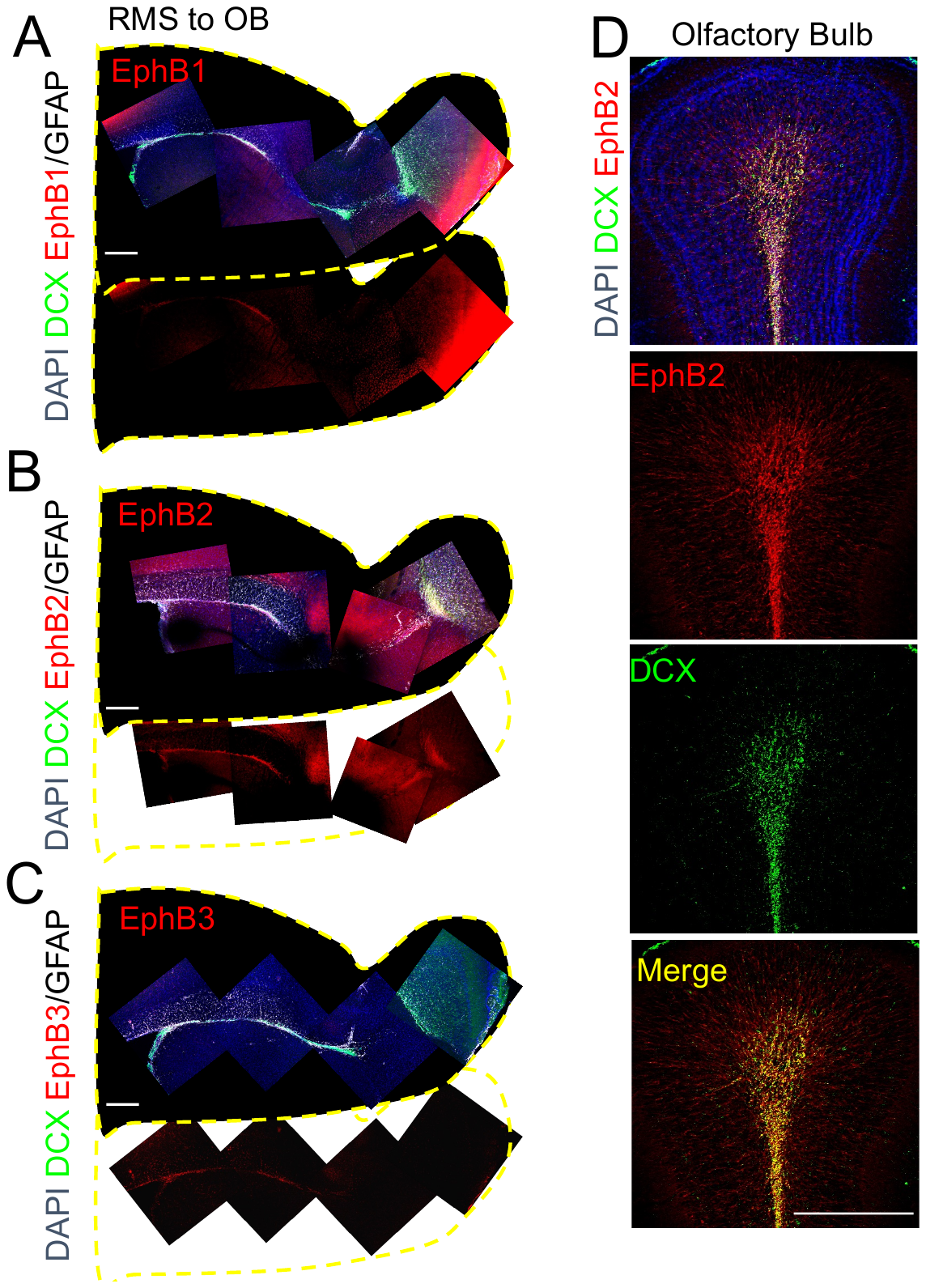

**Supplementary** **Figure 2. EphB expression in the RMS and olfactory bulb.** (**A**) Low magnification composite image (imaged regions shown as squares in area of black of EphB1 immunostaining in the RMS and olfactory bulb. Outline of region in yellow show section. Top panel shows merged image of α-EphB1 (red), DCX (green), and GFAP (white). Lower panel shows α-EphB1 staining alone (red). (**B**) As in **A** but showing α-EphB2 staining. (**C**) As in **A** but showing α-EphB3 staining. (**D**) images of the olfactory bulb stained with α-EphB2 (red), anti-DCX (green), and DAPI. Lower panels show α-EphB2, α-DCX, and EphB2-DCX merged images as labeled. Orientations are labeled in image as Dorsal (D) and Rostral (R). Scal bars=500 μm, in **A**-**C** and **D**.

**Supplementary** **Figure 3. EphB2 expression in the RMS.** (**A**) Adult mouse brain sectioned and labeled for α-EphB2 (red), α-NeuN (Green) and α-GFAP (blue) and DAPI (white). Large panel shows merged image. Small panels show magnified view of inset box. Upper left panel: α-EphB2; upper right panel: α-GFAP; lower left panel: DAPI; lower right panel: α-NeuN. (**B**) Large panel shows merged image of α-EphB2 (red), α-DCX (green), and DAPI (blue) staining. Small panels: α-EphB2 (red), α-DCX (green), DAPI (blue) and merged images (red+green = yellow). All images are from sagittal sections. Orientations are labeled in image as Dorsal (D) and Rostral (R). Scale bars =100 μm, 20 μm.

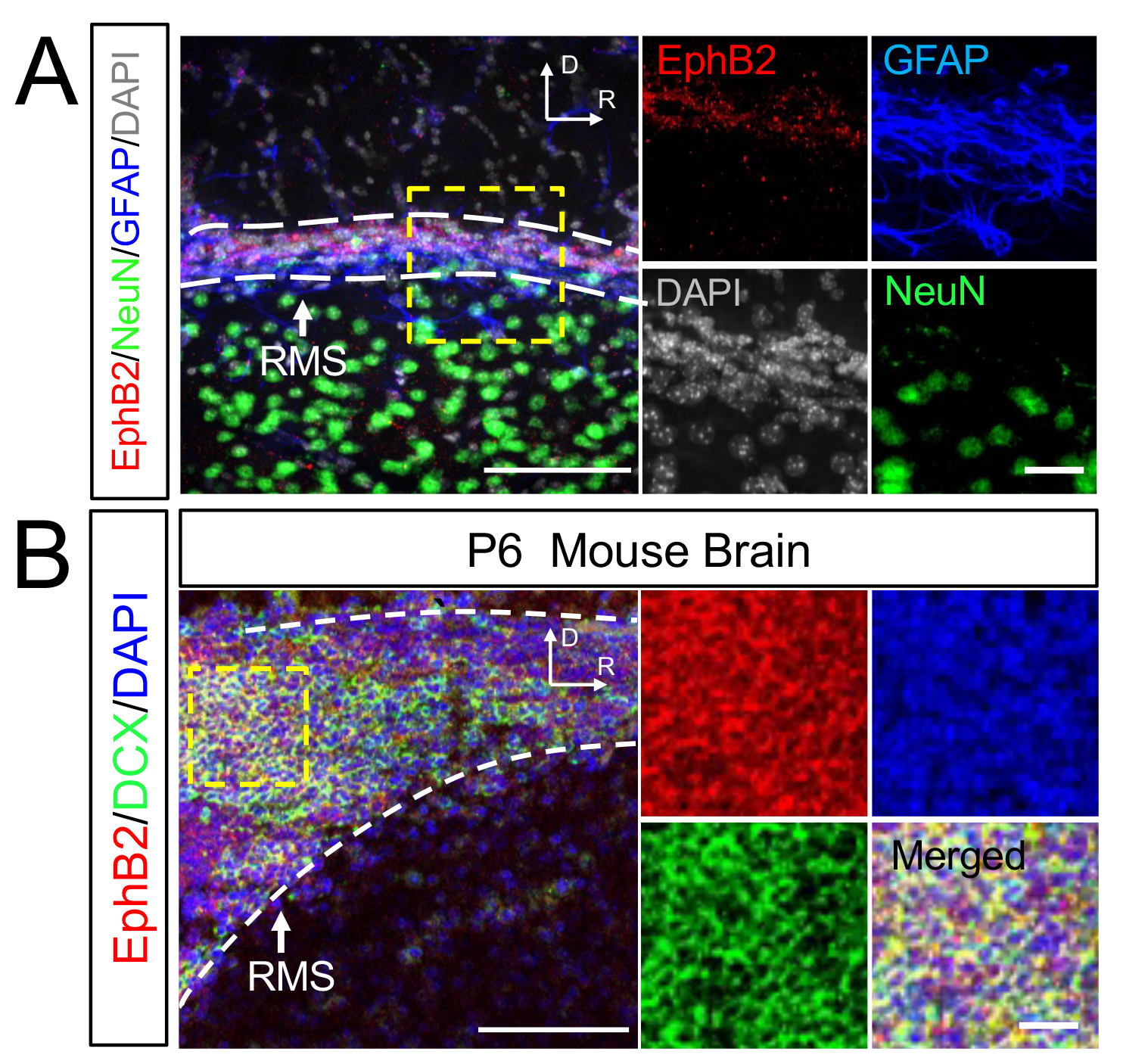

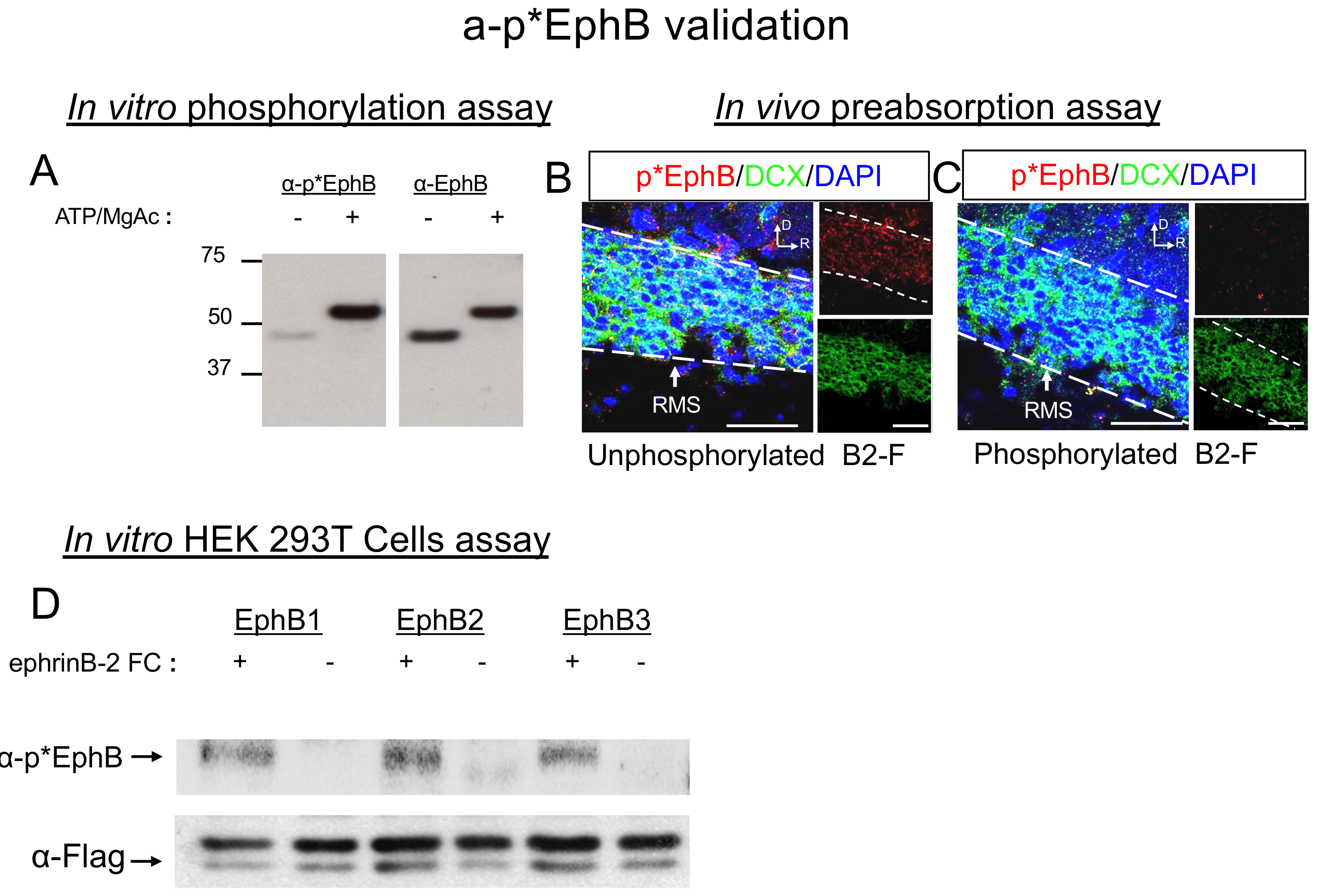

**Supplementary Figure 4. Phospho-EphB (p*EphB) antibody validation.** (**A**) Western blots of EphB2 intracellular domain including the kinase domain incubated with 10 mM MgAc alone (-) or 100 μM ATP and 10 mM MgAc (+) for 30 °C for 30 min. Western blotting with α-p*EphB2 or α-EphB2 showed that intracellular protein fragment of EphB2 is phosphorylated in the presence of ATP and MgAc (left). The total amount of EphB2 was comparable among samples (right). (**B**-**C**) Specificity controls for p*EphB antibody: EphB2 intracellular domains were pre-phosphorylated using the protocol in (A) and incubated with brain sections, and stated for p*EphB2. (**B**) Brain sections were stained for DAPI (blue), α-DCX (green), and α-p*EphB (red, incubated with unphosphorylated EphB2 intracellular domain (B2-F). Large panel shows merged high contrast image with DAPI (blue), α-DCX (green), and α-p*EphB (red). Upper right panel: α-p*EphB; lower right panel: α-DCX. (**C**) Brain selections incubated with phosphorylated EphB2 intracellular domain (B2-F) stained as in **B. (D)** Western blot showing α-p*EphB HEK293T cells transfected with FLAG-tagged EphB1, EphB2 or EphB3 and treated with ephrin-B2-FC to activate EphB kinases. Top Blot shows p*EphB staining. Bottom blott shows FLAG staining. All images are from sagittal sections. Orientations are labeled in image as Dorsal (D) and Rostral (R). Scale bar =50 μm.

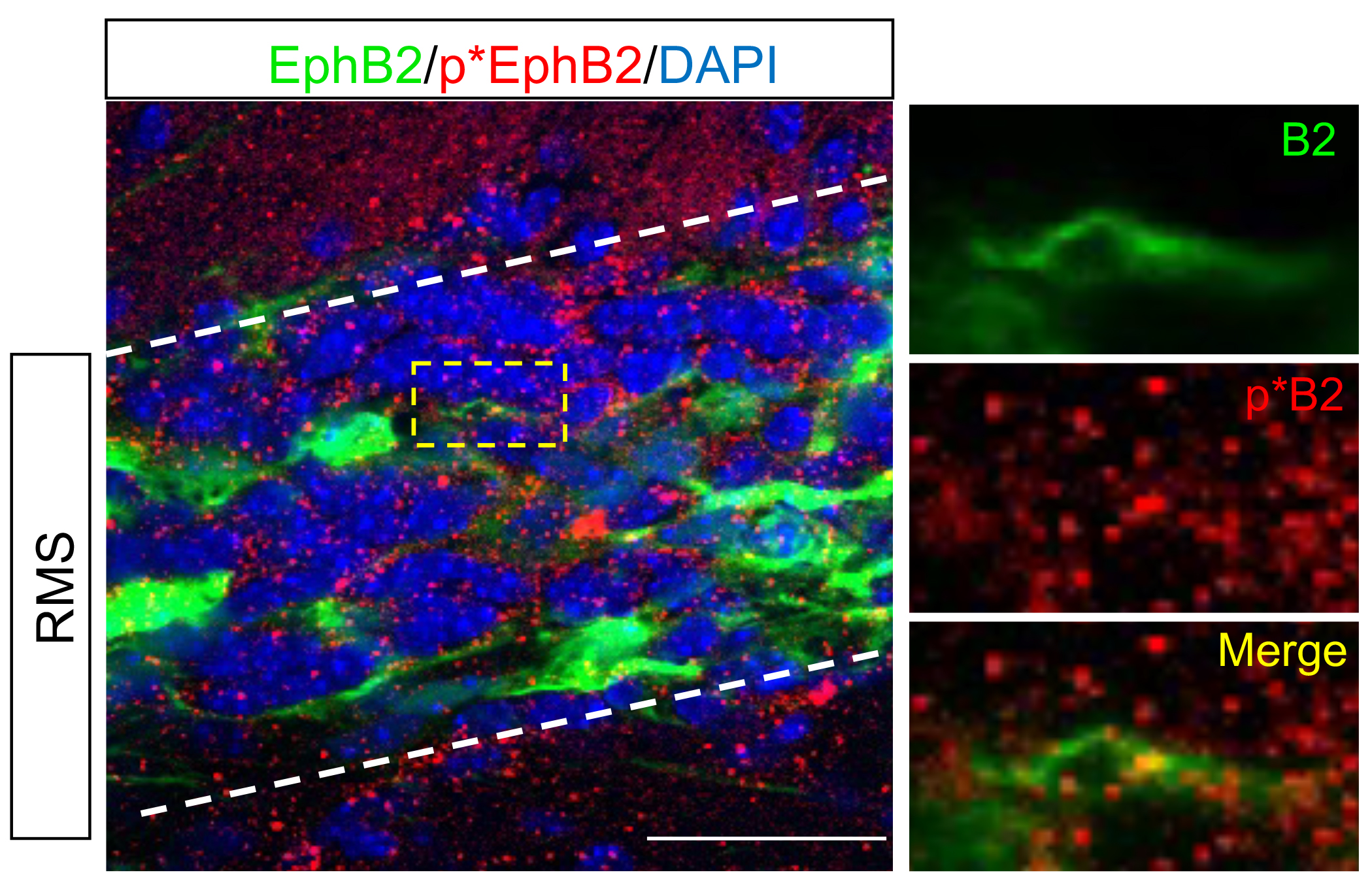

**Supplementary Figure 5. High magnification image of EphB2 and pEphB staining in RMS.** Example of EphB2 positive cell with a migrating profile in the RMS. Large image shows RMS stained for EphB2 (green), pEphB (red), and DAPI (blue). Dashed lines indicate boarder of RMS. Images on the right show magnified area in yellow box. Scale bar = 100um.

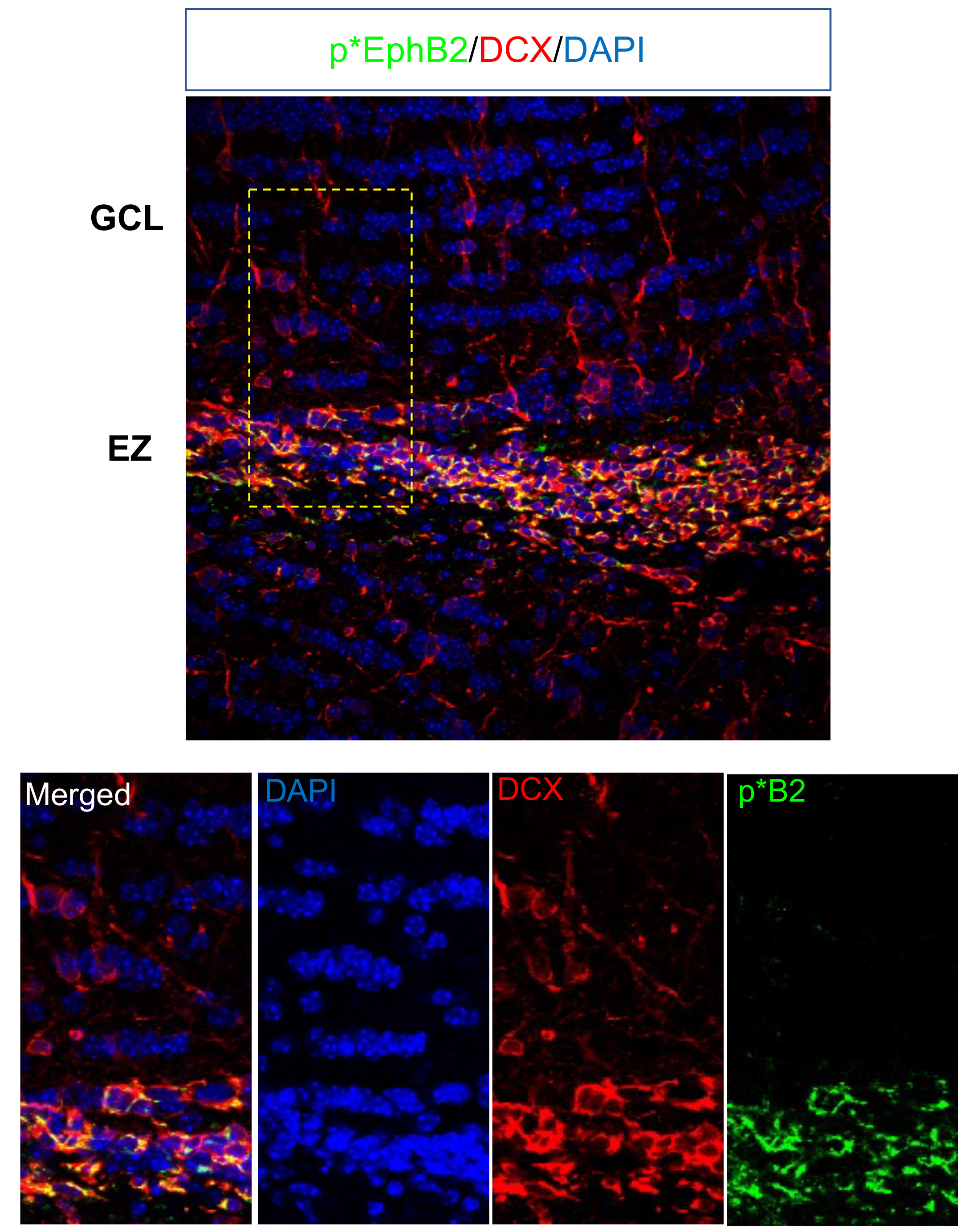

**Supplementary Figure 6. Example of pEphB staining in EZ and GCL of the olfactory bulb.** Top image shows merged image of GCL and EZ stained for pEphB (green), DCX (red) and DAPI (blue). Lower images show enlarged region boxed in yellow dashed line as labled left to right: merged, DAPI, DCX, p*EphB (p*B2).

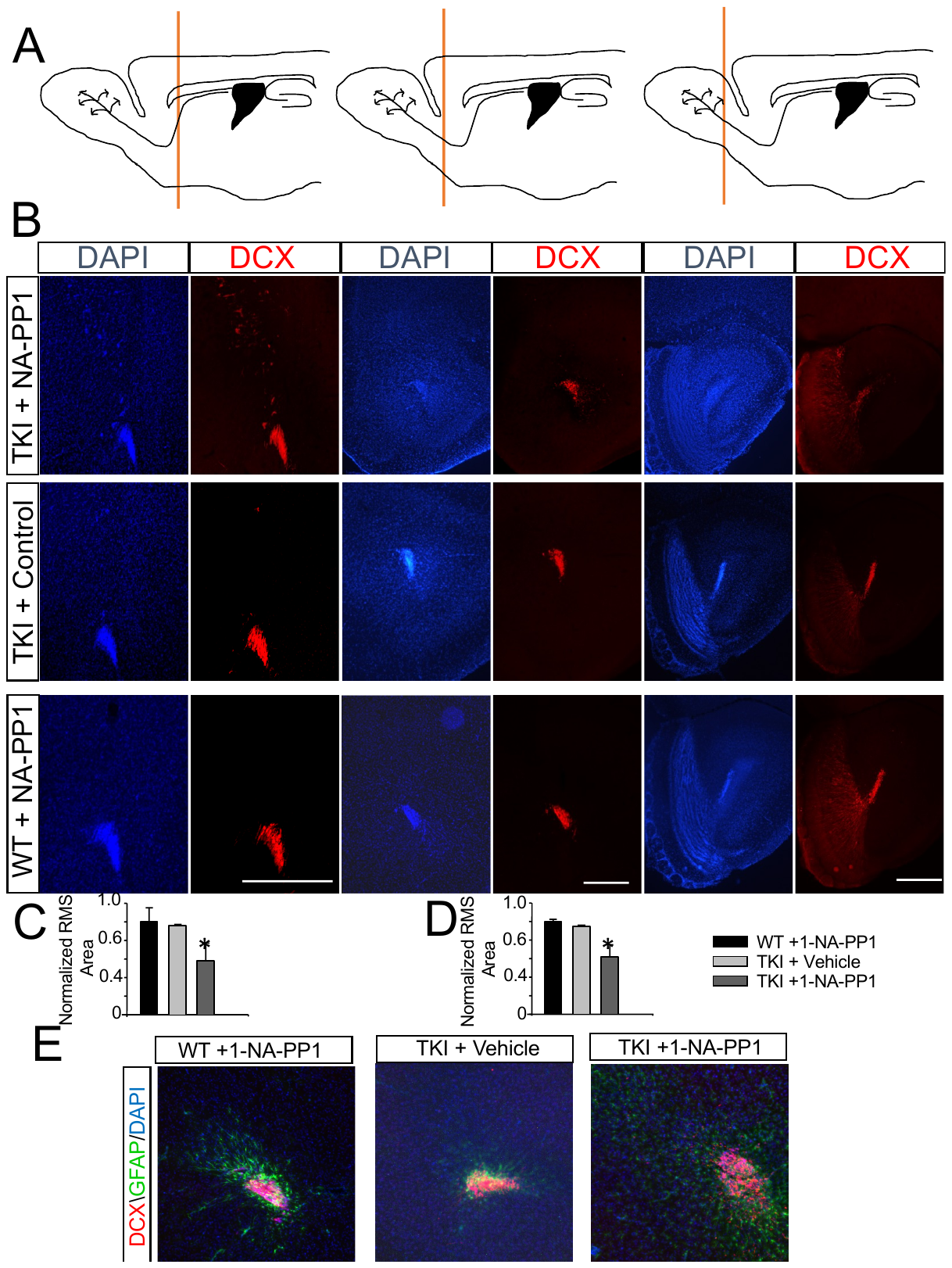

**Supplementary Figure 7: Corronal sections through RMS in TKI mice.** (**A**) Diagrams of the locations of the three different sections shown in (**B**) from caudal to rostral with the red vertical line indicating the approximate position of the sections. (**B**) Example sections from three different locations along the RMS (caudal to rostral). Images are shown DAPI (blue) and DCX (red). Top panel is in TKI mice with control treatment, middle panel shows TKI mice plus NA-PPI to block EphB kinase activity and bottom images show WT mice treated with NA-PPI. (**C-D**) Quantification of effects of NA-PPI on the area of the RMS. (E) Higher magificatin views of coronal sections through the RMS stained for DCX (red), GFAP (green) and DAPI (Blue). Scale bars = 800um (A), 400um (E). * p<0.05

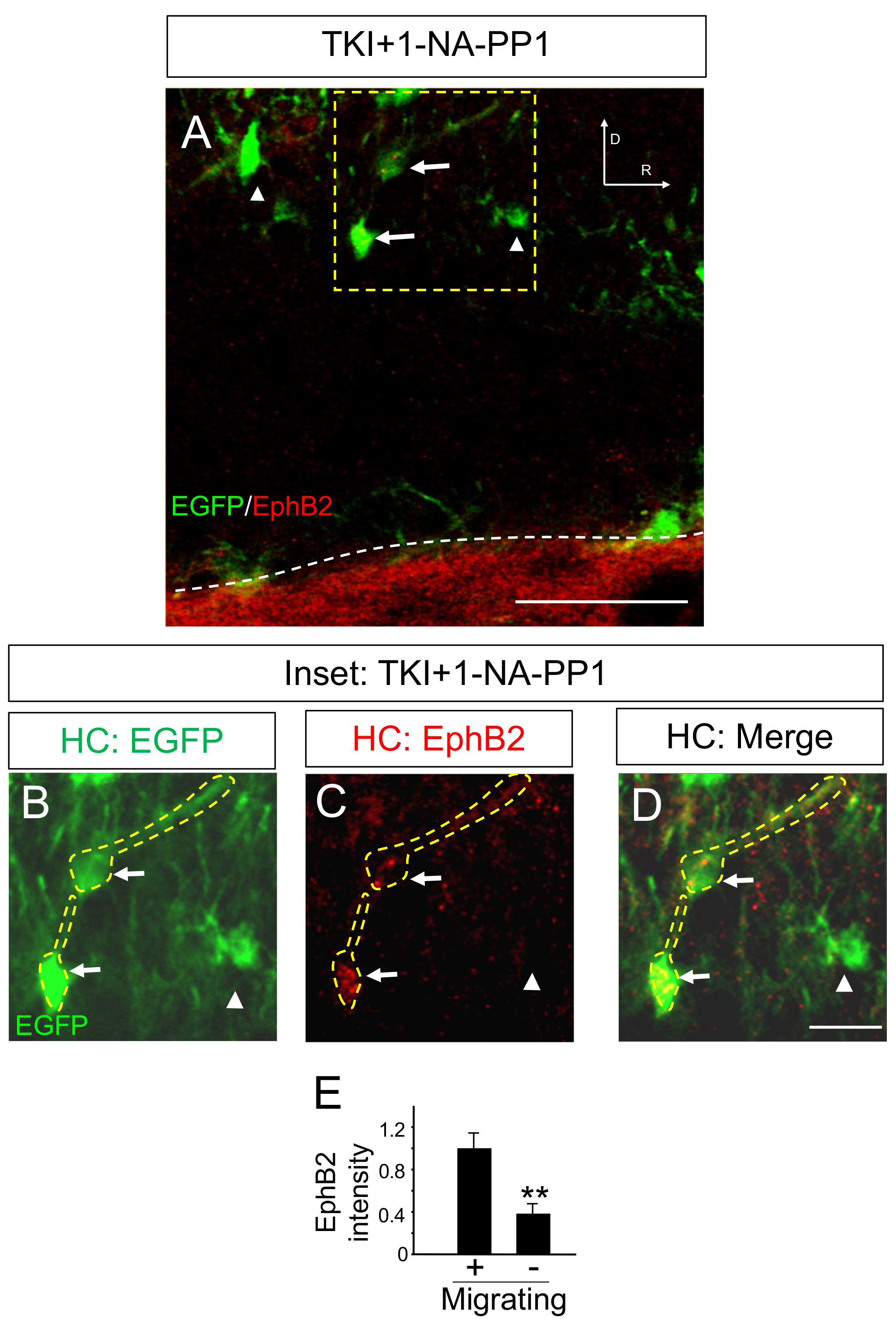

**Supplementary Figure 8. EphB2 staining in EGFP expressing cells following inhibition of EphB kinase activity.** (**A**) Low magnification image of EGFP+ cells labeled by injection of LV transducing EGFP in the lateral ventricle in TKI mice following injection of NA-PP1. RMS is indictated by the dashed line and EphB2 staining (red). Examples of cells with a complex morphology (Arrowhead) and migrating profile (Arrow) are shown. (**B-D**) Higher magnification image of region boxed by dashed yellow line in A. B. Shows EGFP channel (green), C shows EphB2 staining (red), D shows merged image. E. Quantification of normalized EphB2 expression in EGFP+ cells of migrating morphology (arrow in A) and differentiated morphology (arrowhead in A) (n=30 cells/group). n=4 animals per group. All images are from sagittal sections. Scale Bar = 200 μm A, 50 μm (B-D). ** p<0.01, Student’s Test.

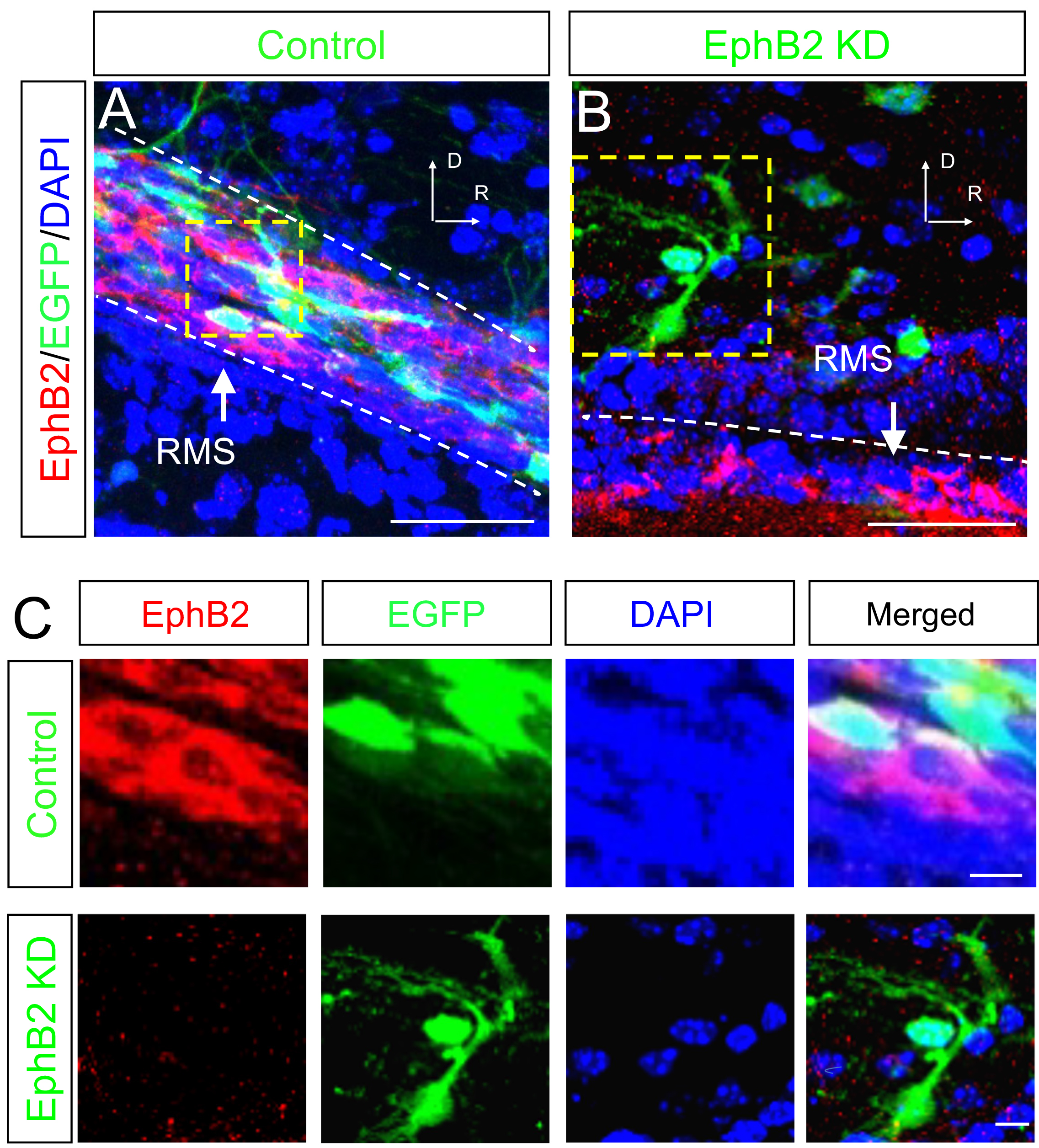

**Supplementary Figure 9. Effects of EphB2 shRNA knockdown in RMS** (**A-B**) Validation of EphB2 shRNA. (**A**) Control RMS injected with control EGFP transducing lentivirus stained for α-EphB2, α-EGFP and DAPI. Large panel shows merged image of EphB2 (red), EGFP (green), and DAPI (blue). Small panels show magnified view of inset box stained as indicated. (**B**) RMS injected lentivirus transducing EphB2 shRNA and EGFP stained for as in **A**. (**C**) Z-projection for EphB2 shRNA transduced EGFP+ and GFAP+ cells shown in Fig. 9**D**. (**D**) Example of NeuN+/EGFP+ EphB2 shRNA transduced cells that are distant from the RMS (RMS is not pictured). All images are from sagittal sections. Scale bars = 50 μm, 10 μm in panel **A**-**B**, and 50 μm, 20 μm in panel **C**­-**D**.

­­­
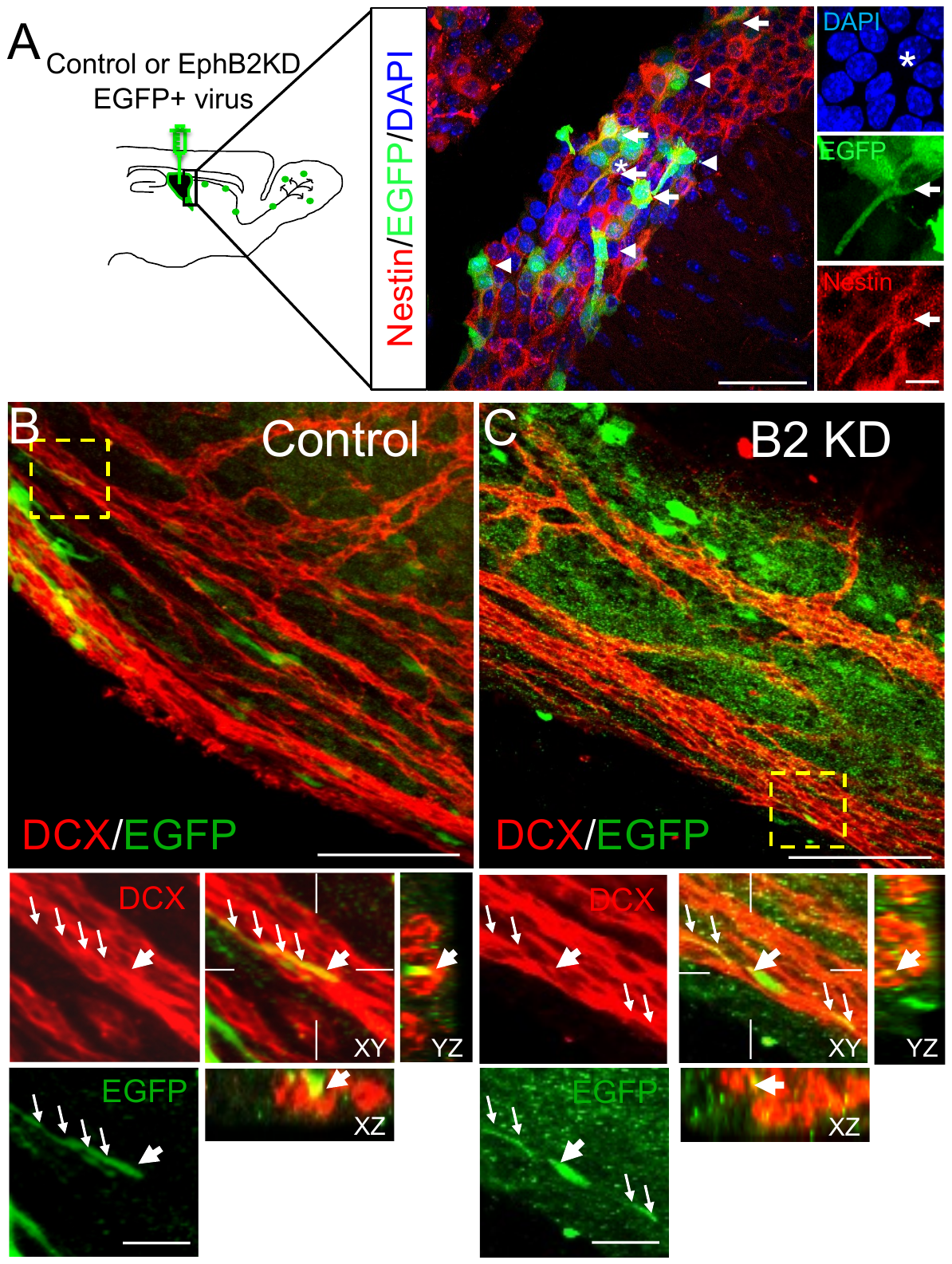

**Supplementary Figure 10. EGFP labeled cells in SVZ and effects of EphB2 knockdown.**

Determination of the effect of EphB2 knockdown in the SVZ. (**A**) Lentiviral injections of EGFP transducing virus labeled nestin+ cells in the SVZ. Lower right panel: α-nestin immunostaining (red), middle right panel: EGFP (green), upper right panel: DAPI (blue), large image is merged. (**B**-**C**) Whole-mount from animals transduced with either control or EphB2 shRNA EGFP lentivirus virus. (**B**) Control injected - large panel shows a merged image of EGFP (green) and DCX (red) labeling. Small panels: upper left: DCX (red); lower left EGFP (green); upper right small merged image with Z projections. Z-projections (xz, yz) are as indicated to show overlap of DCX and EGFP staining (yellow). Arrows indicate position of EGFP labeled neuroblast. (**C**) EphB2 shRNA injected (B2 KD). Panels are as in **B**. Scale bars =50 μm, 10 μm in panel **A**, 100 μm, 20 μm in **B**-**C.**

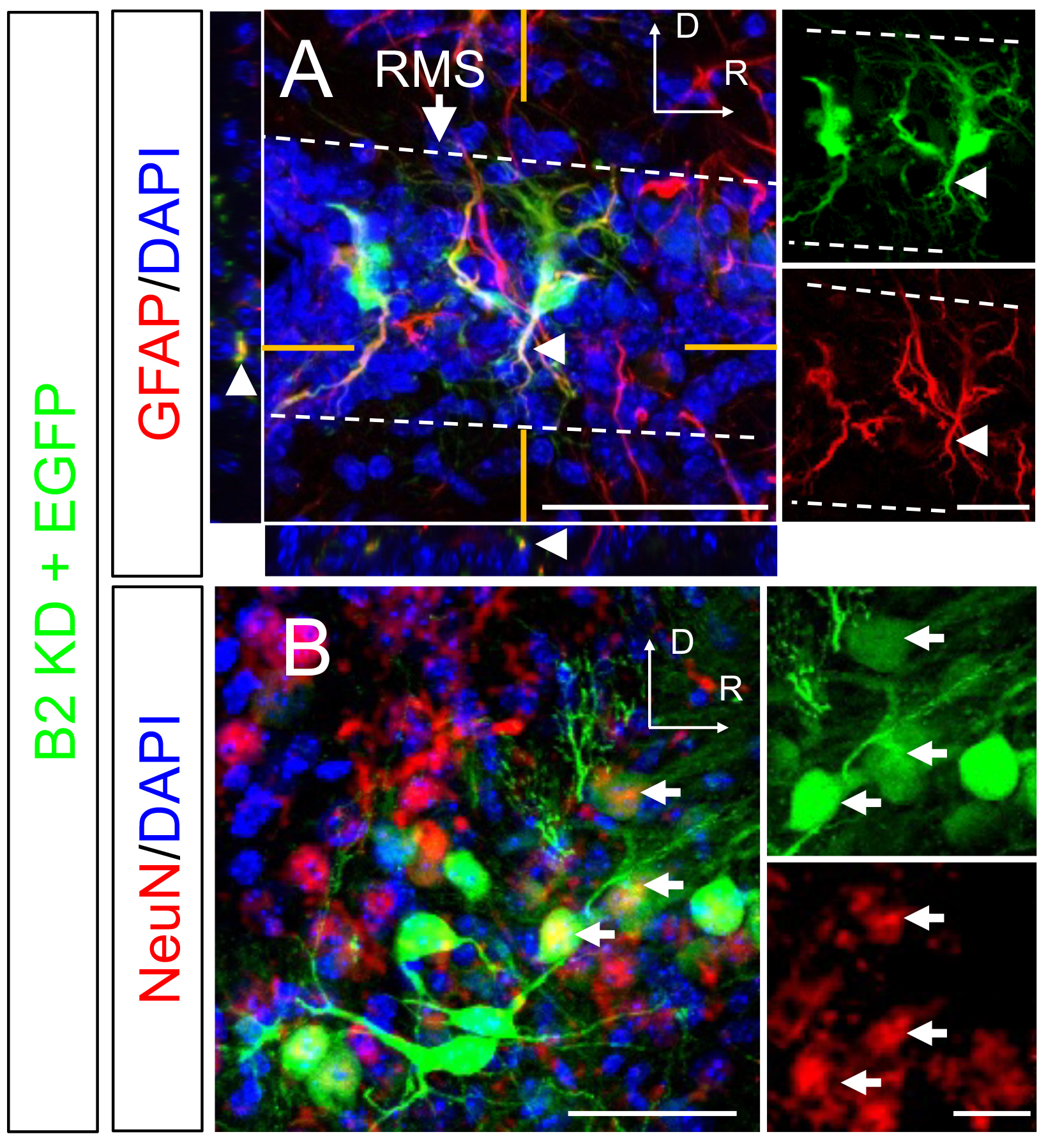

**Supplementary Figure 11. Effects of EphB2 knockdown on neuroblasts.** (**A**) Example of cells transduced with shRNA targeting EphB2 and expressing EGFP in RMS expressing morphology of glial cells, and expressing GFAP (red). Small side images show z-projections of EGFP, GFAP and DAPI staining. Arrow indicates border of RMS, arrow heads indicate location of process shown in the z-projections to the left and bottom. Smaller panels to right show single color images EGFP (top, green) and GFAP (bottom, red). (**B**) EGFP+ cells transduced with injection of LV expressing EGFP and shRNA targting EphB2 in ventricle located distant from the RMS. Section stained for EGFP (green), NeuN (red), and DAPI (blue). Arrows indicate EGFP+ cells that co-stained with NeuN. Panels to the right show EGFP (top, green), NeuN (bottom, red). Dorsal ventral orientiation of images as indicated. Scale bars 200μm (A, B), 20μm slide panels.

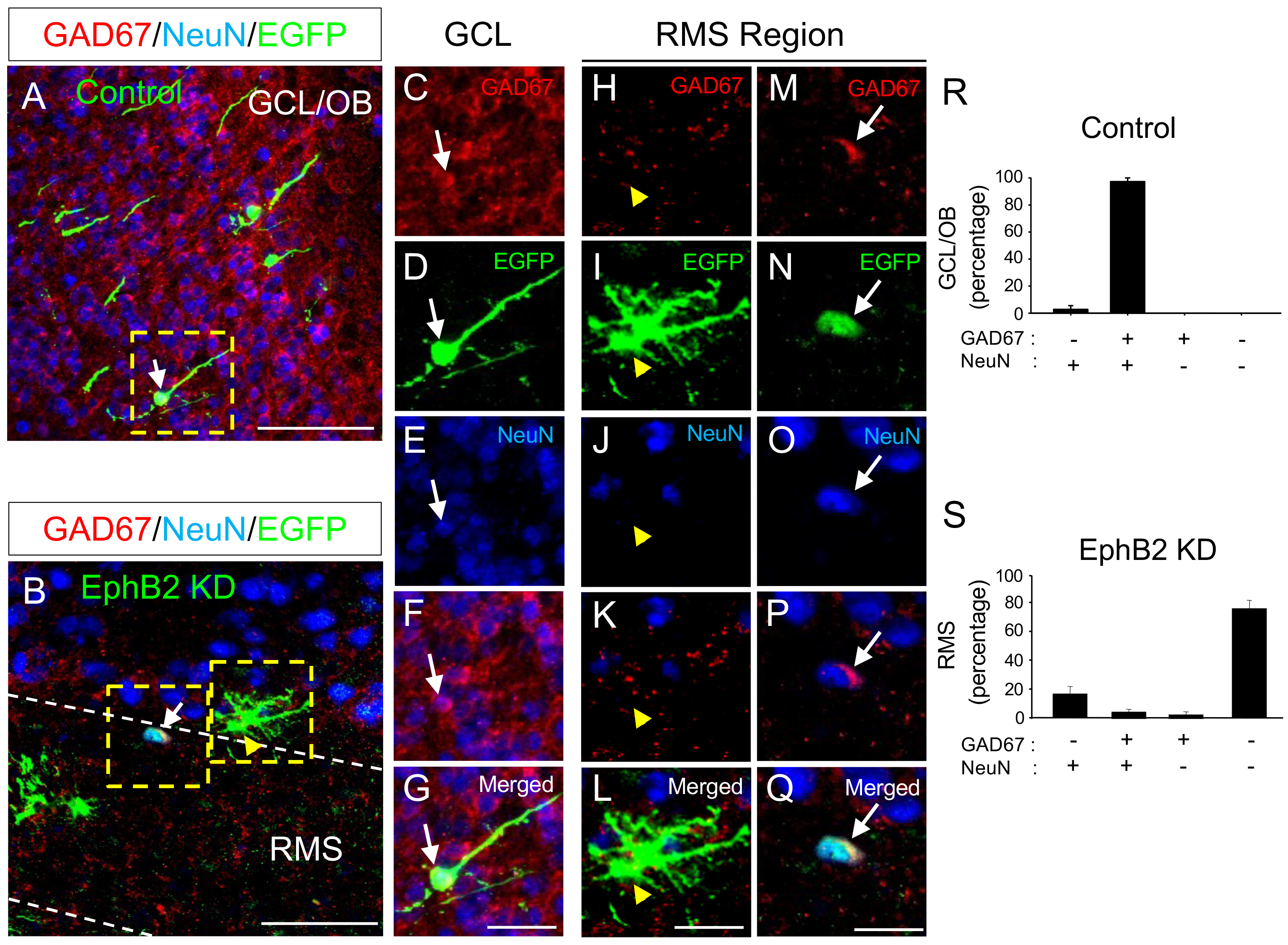

**Supplementary Figure 12. Effects of EphB2 knockdown on neuroblast differentiation.**

(**A**) Control GCL of OB of mice injected control lentivirus transducing EGFP stained for α-GAD67 (red), α-EGFP (green), α-NeuN (blue). (**B**) RMS of mice injected lentivirus transducing EphB2 shRNA and EGFP stained as in a. (**C-G**) Magnified view of GCL inset box of **A showing a** NeuN+/GAD67+/EGFP+ indicated by arrows. Staining as labeled. (**H-L**) Magnified view of inset box (right) in **B** showing a NeuN-/GAD67-/EGFP+ cell in RMS indicated by yellow arrow heads. (**M-G**) Magnified view of inset box (left) in **B** showing a NeuN+/GAD67+/EGFP+ cell in RMS indicated by arrows. (**R**) Quantification of percentage of EGFP+, GAD67+ and/or NeuN+ cells in GCL of OB in control injected brains (n=60 cells). (**S**) Quantification of precentage of EGFP+ cells, GAD67+ and/or NeuN+ in RMS in EphB2 shRNA transduced brains (n= 170 cells). All images are from sagittal sections. Scale bars =100, 20 μm,
